# Secondary motor cortex tracks decision value and supports behavioral flexibility during non-instructed choice

**DOI:** 10.1101/2024.10.29.620931

**Authors:** Elisabete Augusto, Vladimir Kouskoff, Nicolas Chenouard, Margaux Giraudet, Léa Peltier, Aron de Miranda, Alexy Louis, Lucille Alonso, Frédéric Gambino

## Abstract

Optimal decision-making relies on interconnected frontal brain regions, which permit animals to adapt their decisions based on their internal state, experience, and environmental context. Among them, the secondary motor cortex (M2) shows earlier decision-related activity required for sensory-guided action selection. However, the role of M2 in adaptive decision-making in the absence of instructive sensory cues remains unclear. Under such conditions, action-selection relies on abstract representations of actions and their values. Using *in vivo* microscopy and modeling, we showed that M2 neurons in mice exhibited persistent activity encoding decision values (DV) predicting the probability of action-selection during a non-cue-guided lever task. This was confirmed by the reduced reversal performance upon M2 optogenetic inhibition prior to action-selection. Furthermore, updates in DV determined the rate at which learning is reversed. Together, these results provide strong evidence of the use of DV by M2 to adapt choice in the absence of instructive sensory cues.

**Declaration of interests:** The authors declare no competing financial interests.

## INTRODUCTION

Cognitive flexibility is crucial for learning and survival. It involves the capacity to rapidly adapt behavioral strategies and actions in an ever-changing environment in order to optimize valuable outcomes. It encompasses a wide range of different brain processes, such as the ability to acquire novel response strategies and switch between them, or to reverse learning by disengaging from previously learned associations (Brown & Tait, 2014; Soltani & Izquierdo, 2019). The neural mechanisms of cognitive flexibility have been predominantly investigated in the context of perceptual decision-making (Banerjee et al., 2020; Floresco & Ghods-Sharifi, 2007; Haddon & Killcross, 2006; Rich & Shapiro, 2009; Siniscalchi et al., 2016). These studies employed a range of sensory modalities to instruct the action, emphasizing the role of arbitrary sensorimotor association. Nevertheless, individuals must often resolve conflicts between actions while the environment remains apparently unchanged due to the absence or ambiguity of sensory cues. In such cases, the subjective values (*Q*) that individuals assign to potential actions become the primary determinants of their choices (X. J. Wang, 2008). These values are constantly updated by the animal’s ongoing experience of action-outcome contingencies and by its internal state. These values are known to guide choices and require animals to compare the value of each option by computing the difference (Δ*Q*) between them (Dayan & Daw, 2008; Krajbich et al., 2010; Maia & Frank, 2011; Morris et al., 2014; Sutton & Barto, 2018; X. J. Wang, 2008). These differences Δ*Q*, defined here as the decision-values (DV), are not directly observable but can be inferred from reinforcement learning (RL) models, providing comprehensive algorithms to explain behavioral strategies. RL models have greatly inspired decision-making experiments, but the neural circuits that govern them in the absence of predictive external stimuli are not yet fully understood.

Previous studies have revealed neuronal signatures related to DV in many brain areas, particularly in the frontal cortex (Bari et al., 2019; Cazettes et al., 2023; Hattori et al., 2019; Hattori & Komiyama, 2022; Sul et al., 2011; Tsutsui et al., 2016). For instance, DV has been shown to be encoded in the medial prefrontal cortex (mPFC) (Bari et al., 2019; Murakami et al., 2017), but this region does not appear to play a role in reversal learning (Bissonette et al., 2008; Floresco et al., 2008). In contrast, damage to the orbitofrontal cortex (OFC) in rodents appears to be critical for reversals based on sensory stimuli (Bohn et al., 2003; Burke et al., 2009; Schoenbaum et al., 2002) although the OFC does not seem to encode a specific option at the time of choice (Sul et al., 2010). Instead, OFC infers the value of each available option, which can be used elsewhere in the brain to decide the next action (Balewski et al., 2022; Padoa-Schioppa, 2013; Padoa-Schioppa & Assad, 2006; Rich & Wallis, 2016; Wilson et al., 2014). As a result, DV is weakly represented in the OFC (Hattori et al., 2019; Sul et al., 2010). Together, these observations have underscored the difficulty to identify which region compares actions’ values by encoding DV, particularly in the absence of perceptual information.

The secondary motor cortex (M2) has recently attracted significant attention and emerged as a critical structure in rodents, orchestrating motor planning and spontaneous action initiation (Barthas & Kwan, 2017; Ebbesen et al., 2018; Ebbesen & Brecht, 2017). The M2 is connected to brain structures encoding action values such as the mPFC and the OFC (J. H. Yang & Kwan, 2021), exhibits early decision-related activity (Cazettes et al., 2023; Hattori et al., 2019; Hattori & Komiyama, 2022; Sul et al., 2011), and is essential for actions guided by sensory antecedents (Barthas & Kwan, 2017; Ebbesen et al., 2018; Siniscalchi et al., 2016). This structure contributes also significantly to learning and reversal learning (Gremel & Costa, 2013). Therefore, we hypothesized that during a learning and reversal learning task in a non-instructed environment, the M2 builds abstract representations and selects the optimal action between two options by computing, tracking and keeping online the difference between their respective values. This process may provide an important internally-generated representation of the most valuable action, despite an ambiguous context, which is critical for optimal decision-making. To address this hypothesis, we designed a simple two-options behavioral task in which head-restrained mice first acquire and then reverse reinforced self-initiated lever-pressing preferences, without any external cues indicating the rewarded action. The reinforced choice is driven by the difference in option values (*i.e.* DV), which is updated by choice history. During our task, mice actively use their forepaws to make choices, which may preferentially engage frontal cortical areas (Gilad et al., 2018). Accordingly, using two-photon calcium imaging of M2 neuronal population activity combined with behavioral modeling and optogenetics, we showed that the M2 encodes information about DV through persistent population activity, which could serve as a deterministic signal later used to dictate the probability of taking each action. We found that the persistent coding varies slowly from trial to trial and reflects how DV is updated after each action-outcome pair is observed. This, in turn, determines the rate at which learning occurs and is reversed when the reward contingency changes unexpectedly.

## RESULTS

### Reversal of learned self-initiated choice in head-fixed mice

We used a modified self-initiated, two-choice lever-pressing “bandit” task with deterministic reward schedules to test the ability of head-fixed mice to learn and reverse choice behavior. Our “reversal task” does not necessitate the active storage of an external instructive stimulus in memory, and involved three consecutive phases: training, learning and reversal (**Fig.1A; SupFig.1**). First, naive water-restricted mice were trained to press two levers for reward with the corresponding left or right forepaw. The mice self-initiated a unique choice on each trial by pressing one of two adjacent levers (left or right), each of which was then retracted and reinforced by a drop of water from a single spout. Therefore, during training mice learn to associate reward with lever press, and the difference in lever choices enabled us to identify their preference for the left or right lever (**SupFig.1B**). Pressing this preferred lever was subsequently designated as ‘action 1’. Once the mice successfully reached ∼100 presses per session (≤ 1 hour), they began the learning cycle on the next daily session. During learning, the preferred action 1 was no longer rewarded (miss trials), only the press of the opposite lever of their initial bias (action 2, left or right) was rewarded (hit trials; **Fig.1A, B; SupFig.1C, D**). Thus, the choice probability of action 2 increased, while that of action 1 decreased (**Fig.1B, C**). Pressing both levers never led to rewards and occurred only occasionally. Once mice had learned the rule policy of the task (>75% of rewarded choices for 3 consecutive sessions), the reversal learning cycle began, during which the reward contingencies were unexpectedly reversed (i.e. without any instruction), resulting in an immediate drop in rewarded lever presses. During both learning and reversal learning, the mice did not update their behavioral strategy rapidly, requiring a similar number of sessions to achieve expertise (learning: 19.7 sessions ± 2.6, n= 25 mice; reversal learning: 20.1 sessions ± 2.6; n = 23 mice; **Fig.1C**). This suggests that mice systematically use a trial-and-error approach to update the values linked with each action whenever the reward schedule changes (*i.e.* between training and learning or between learning and reversal learning). This process would devalue the action that was preferred during training or reinforced during learning, while gradually increasing the value of the action that was newly rewarded during learning or reversal learning, respectively. On the other hand, the mice stopped pressing the levers when water was provided *ad libitum*, demonstrating the use of a goal-directed strategy with a rapid disengagement from the task when the outcome is devaluated as satiety increased (Bouton & Balleine, 2019) (**Fig.1D**). Overall, our task shows that head-fixed mice adapt their behavior by disengaging from a previously acquired rule to select a newly reinforced option in a trial-and-error goal-directed manner.

**Figure 1:**
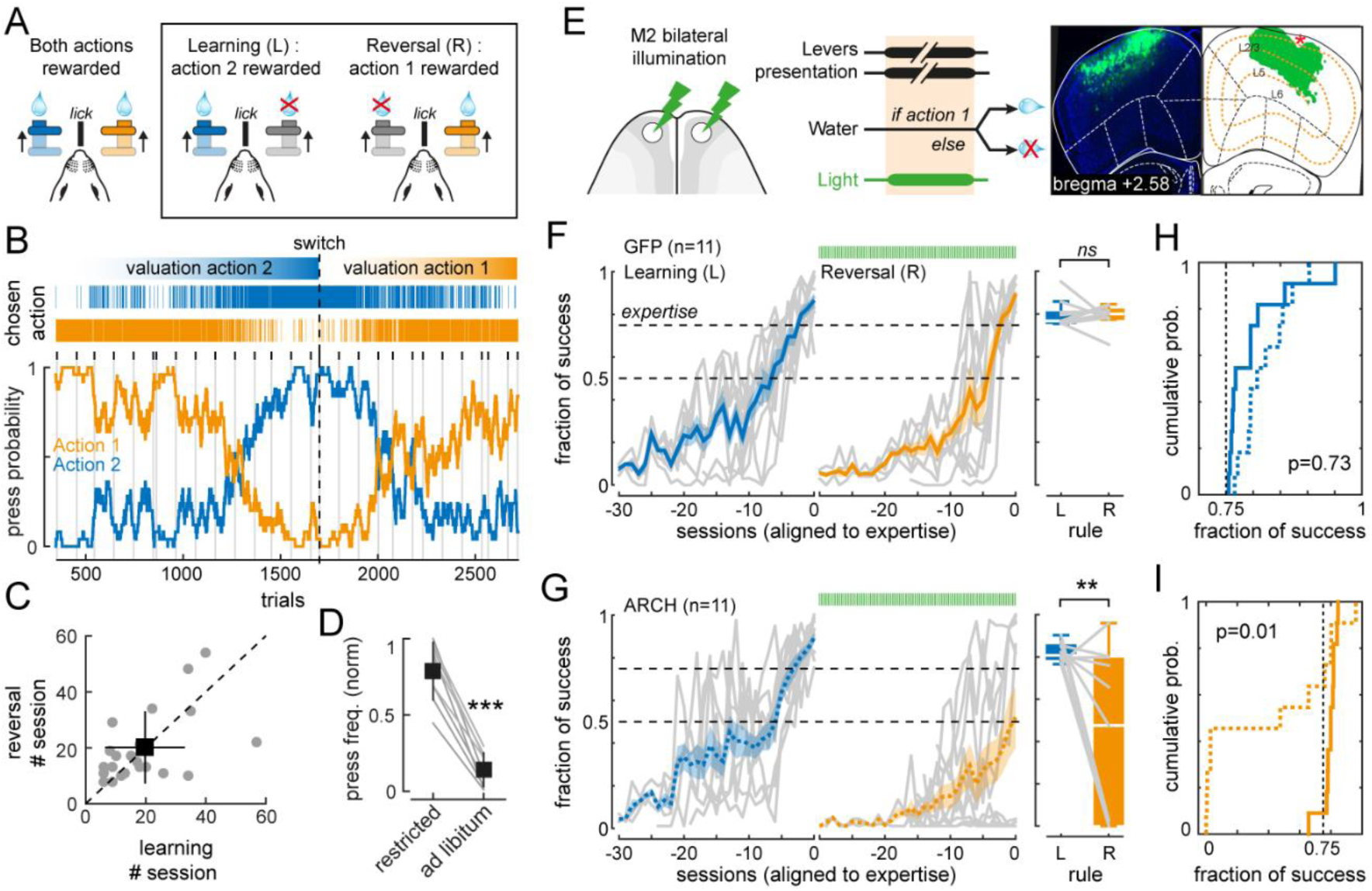
M2-dependent reversal learning task. **A)** Schematic representation of the task during which the mouse decides to press the right (action 1) or left lever (action 2) that retracts afterwards. The task has different phases (training: *left panel*: training trial with two levers presented at the same time that yield the same reward after press; *middle panel*: learning (L) trial with two levers presented at the same time that yield different probabilities of reward after press (100% for action 2 and 0% for action 1); *right panel*: reversal (R) trial with two levers presented at the same time that yield different probabilities of reward after press and opposite of learning trial (100% for action 1 and 0% for action 2). **B)** Learning cycle followed by reversal learning cycle from one mouse, showing the probability of each action across trials. Dashed line marks the switch between learning and reversal. Orange line is the probability of action 1, blue line is the probability of action 2. Grey lines and black ticks show the delimitation of sessions. Top lines reveal chosen action in each trial: blue lines for each time action 2 was chosen, orange lines for each time action 1 was chosen. **C)** Correlation between number of sessions during learning and reversal learning, each point corresponds to one mouse. Black square represents the mean and error bars are sd. **D)** Frequency of lever pressing in water deprived mice (restricted) and in non-water-deprived mice (*ad libitum*). Each grey line represents one mouse. Black squares represent the mean and Error bars are sem. **E)** *Left panel*: schematic representation of M2 bilateral illumination with yellow light. *Middle panel*: schematic representation of the timing of the events within the trial, namely top) presentation of both levers simultaneously, middle) reward or absence of it, depending on the action that was previously executed, action 1 or action 2 respectively, bottom) M2 illumination simultaneously with the presentation of the levers. *Right panel*: histological representation of the coordinates of ARCH viral infection (in green) and optical fiber implantation (red star). **F)** Effect of M2 illumination in GFP-expressing mice. *Left panel*: mean of learning (blue line) and reversal learning (orange line) curves aligned to expertise, depicted as the fraction of successes (action that gives rise to a reward) in the different sessions. Each grey line represents the curve of one mouse. Green bars at the top represents M2 illumination that happens during levers presentation, only on reversal cycle. Dashed line at 0.5 and 0.75 of the fractions of success representing respectively random choices and expertise level. *Right panel*: average of the fraction of successes of the 3 last sessions represented on the left panel (fraction of successes when mice are experts). Blue box is from learning cycle, orange box is from reversal cycle and grey lines represent the behavior of each mouse. Boxplots represent mean and interquartile range. Error bars are sd. **G)** Effect of M2 photoinhibition in ARCH-expressing mice. Similar to f. **H)** Cumulative probability of successes from the 3 last sessions of the learning cycle (blue lines on panels f. and g.) from GFP-expressing mice (filled blue line) and ARCH-expressing mice (dashed blue line). **I)** Similar to h but for the reversal learning cycle.

### M2 neuronal activity before action is required for behavioral flexibility but not movement execution

We tested whether bilateral optogenetic silencing of the M2 during the putative decision period, *i.e.* prior to lever-pressing, impaired reversal learning (**Fig.1E**, left). We induced the expression of the light-activated proton pump *archaerhodopsin* (AAV9.CAG.ArchT.GFP, n=11 ARCH mice) or GFP for controls (AAV9.CamKII.eGFP, n=11 GFP mice), bilaterally in M2 layer 2/3 excitatory pyramidal neurons (L2/3-PNs) and implanted optical fibers in the superficial layer 1 to inhibit preferentially their activity (**Fig.1E**, right). This strategy was previously used successfully to inhibit neuronal activity in M2, and impaired associative learning (Aime et al., 2020). Both groups of mice (GFP and ARCH) trained and learned the task (**Fig.1F, G**), achieving a similar level of expertise in the absence of M2 illumination (GFP: fraction of rewarded trials = 0.8314 ± 0.0623; n = 11; ARCH: 0.8432 ± 0.0555; n = 11; p=0.73; **Fig.1H**). During reversal learning, M2 illumination was synchronized with lever presentation, that is before the action was taken and choice revealed (**Fig.1E**). Unlike GFP mice, whose performance was similar to that of the learning phase (learning: 0.8314 ± 0.0623; reversal: 0.8336 ± 0.0251; n=11; p=0.927; **Fig.1F**), ARCH mice failed, on average, to achieve the same performance (learning: 0.8432 ± 0.0555; reversal: 0.4111 ± 0.3978; n=11; p=0.006; **Fig.1G**). Therefore, ARCH mice had a significantly lower success rate than GFP mice after reversal (GFP: 0.8336 ± 0.0251; n = 11; ARCH: 0.4111 ± 0.3978; n = 11; p=0.01; **Fig.1I**), indicating that neuronal activity in M2 before the action was taken and choice revealed is required to reverse a previously acquired rule.

Neural activity in M2 may initiate complex movements, including grasping and lever pressing (Kawai et al., 2015; Tombaz et al., 2020; W. Yang et al., 2023). Consequently, we quantified motor parameters related to forepaw lever pressing to test whether the decreased behavioral performance during reversal learning was attributable to a disruption of movement-related neuronal activity in M2 (**Fig.2**). First, we calculated the average delay to initiate lever press for all rewarded (hit) and unrewarded (miss) trials, independently of their position in the learning and reversal phases. In contrast to the success rate, we found, on average, no detectable differences in delay between GFP and ARCH mice (**Fig.2A, B**). Notably, the delays did not differ between hit and miss trials, indicating that head-fixed mice can use both paws equally well during the learning and reversal conditions (GFP hit learning: 2.453 ± 1.1082; hit reversal: 2.8078 ± 1.2863; n=11; p=0.288; ARCH hit learning: 2.4565 ± 1.3152; hit reversal: 1.9234 ± 1.1058; n=11; p=0.582; GFP miss learning: 2.6039 ± 0.8794; miss reversal: 2.5447 ± 0.8477; n=11; p=0.751; ARCH miss learning: 2.285 ± 1.0373; miss reversal: 2.7735 ± 1.29965; n=11; p=0.417). Although not being used for making a choice, we also quantified tongue motor output by calculating the mean lick rates and found no effect of M2 photo-inhibition between genotypes (p=0.08; **SupFig.2**). To better quantify motor output, we used advanced machine vision algorithms (Mathis et al., 2018) to track forepaw trajectory during lever press (**Fig.2C**). We observed that lever-press delays covary with traveled distance in both groups, the variance of which thus reflects the consistency of forepaw trajectory (GFP: r² = 0.649; p<0.0001; ARCH: r² = 0.836; p<0.0001; **Fig.2C**). The variance associated to the rewarded action (hit trials) suddenly increased after reversal (**Fig.2D**). This evolved over trials while choice was refined on similar timescales between GFP and ARCH mice, which were thus undistinguishable before and after M2 illumination (normalized variance; GFP; learning: 0.84 ± 0.16; reversal: 1.08 ± 0.09; n=11; ARCH; learning: 0.98 ± 0.08; reversal: 1.11 ± 0.08; n=11; p=0.84; **Fig.2E**). Consistent with previous studies (Kawai et al., 2015; Siniscalchi et al., 2016), our data demonstrate that inactivating the M2 before and during lever-pressing has no significant impact on motor execution.

**Figure 2:**
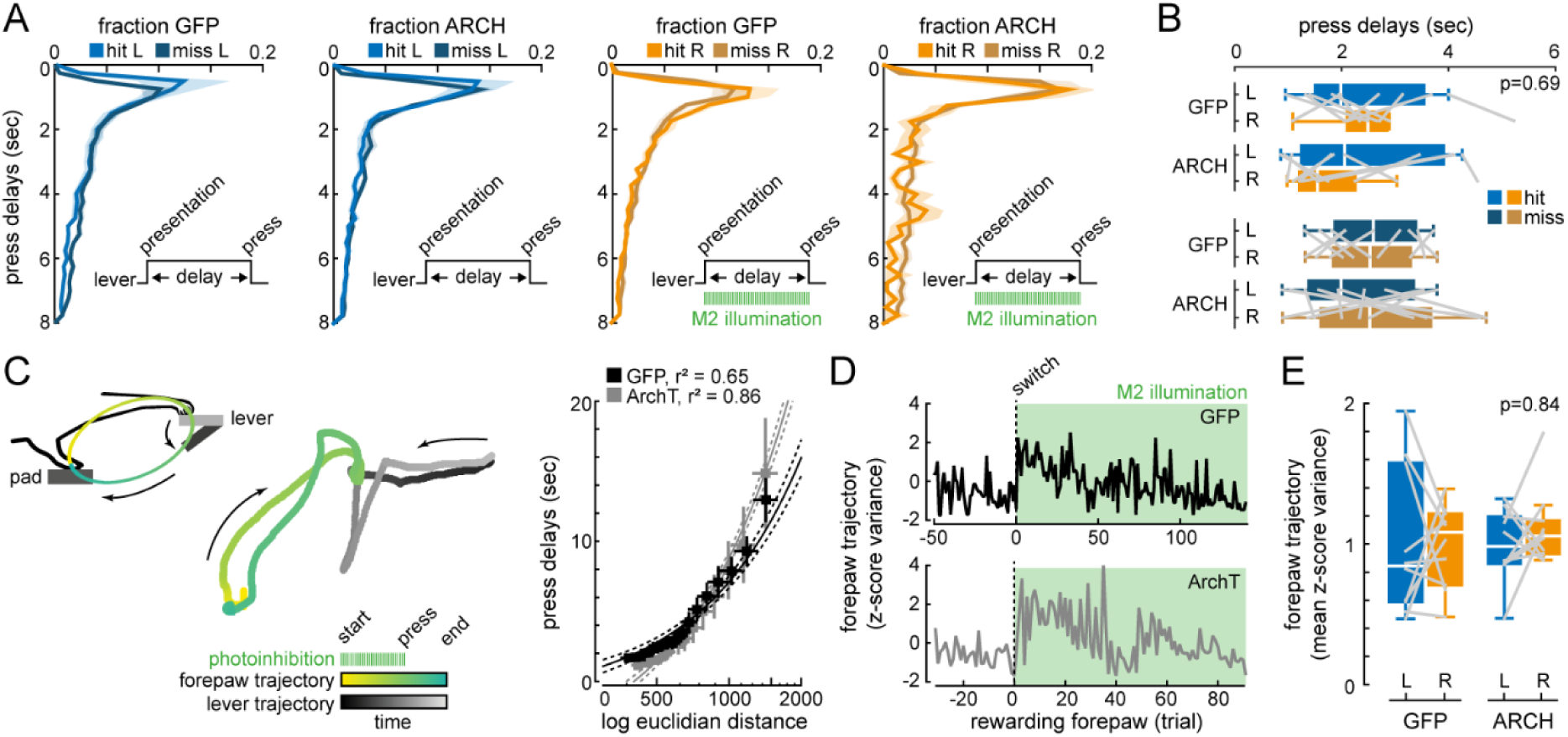
M2 photoinhibition does not impair lever press kinematic. A) Distribution of the press delay (sec), which is the time between levers presentation and lever press, during learning cycle (L, blue graphs) and reversal cycle (R, orange graphs), in GFP- and ARCH-expressing mice. Reversal cycle had light ON during lever presentation. Hit for rewarded trials and miss for unrewarded trials. B) Average of the press delays (sec) in GFP- and ARCH-expressing mice during learning (L, blue graphs) and reversal with M2 bilateral illumination (R, orange graphs), for hit (lighter colors) and miss (darker colors) trials. Boxplots represent mean and interquartile range. Error bars are sd. C) Trajectory of the forepaw during a lever press of the reversal cycle with M2 photoinhibition. Left panel: schematic representation of the movement of the forepaw during a lever press. Middle panel: trajectory of the forepaw of one mouse in one trial (green trajectory) and the trajectory of the lever (grey trajectory). Right panel: correlation between the press delay and the log of the total Euclidian distance between all consecutive time points of the forepaw trajectory, in GFP- (black squares) and ARCH-expressing (grey squares) mice. Each square represents the mean (+/- sd) of distance and delay (bin of 20 ms). D) z-score variance of the forepaw trajectories in hit trials, during the last 50 trials of the learning cycle and the first 100 trials of the reversal cycle (green background). Dashed line marks the switch between learning and reversal cycles. E) Average of the z-score variance of the forepaw trajectories in hit trials, in GFP- and ARCH-expressing mice during the learning (L) and reversal (R) cycles. Boxplots represent mean and interquartile range.

### Reinforcement learning model and optogenetic inhibition uncover decision-value in M2 during reversal task

Learning and reversal learning in rodent are classically described by simple task parameters (e.g., fraction of success or lick rate). To obtain a richer picture, we fitted correct current choice behavior with a logistic regression model to quantify the influence of past choices on the reward rate (3 predictor types: past rewarded choices, past unrewarded choices, reward-independent choices; 31 parameters in total; **SupFig.3A**) (Methods) (Hattori et al., 2019; Katahira et al., 2015). For the predictor “past rewarded choices”, positive regression coefficients that were above chance level decreased as a function of past rewards (**SupFig.3B**), and predicted correct choice patterns with a significantly better accuracy than a simple constant bias parameter (31 parameters-history model: accuracy = 71.9609 ± 8.937 %, n=16; single constant bias model: 58.6446 ± 7.7393 %, n=16; p=0.00009; 2-fold cross-validation) (**SupFig.3C**), suggesting that mice integrate reward history to achieve expertise.

**Figure 3:**
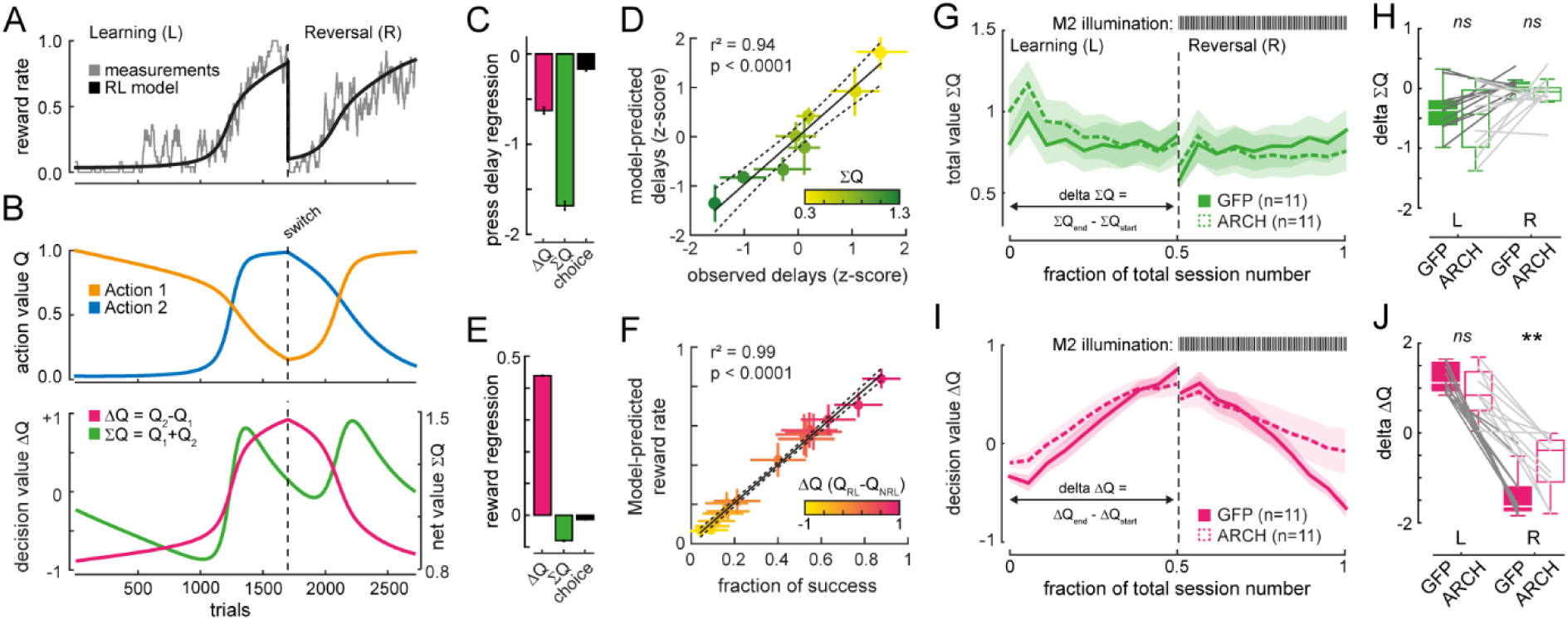
Reinforcement learning model predicts behavior and reveals decision-values. **A)** Reinforcement learning model (black line) fits reward rate (grey line) during learning (before switch) and during reversal learning (after switch). Dashed line marks the switch. **B)** *Top panel*: reinforcement learning model reveals values (Q) associated to action 1 (Q_1_, orange line) and to action 2 (Q_2_, blue line), throughout learning (before switch) and reversal learning (after switch). *Bottom panel*: reinforcement learning model allows to calculate the difference between decision values (ΔQ, pink line) and the sum of decision values (ΣQ, green line), throughout learning and reversal learning. **C)** Regression weight of model parameters predicting press delay. **D)** Correlation between the model-predicted press delays and the observed delays, based on ΣQ. Each circle represents the mean (+/- sd) of binned predicted and observed delays (bin 250 ms). **E)** Regression weight of model parameters predicting reward rate. **F)** Correlation between the model-predicted reward rate and the observed fraction of success, based on ΔQ. Each circle represents the mean (+/- sd) of binned predicted reward rate and observed fraction of success (+/- sd). **G)** Total values (∑Q) variation during learning and reversal learning (with M2 bilateral illumination), separated by the black dashed line, in GFP- (full green line) and ARCH-expressing (dashed green line) mice. Shaded areas, sem. **H)** Delta ΣQ of the learning cycle (without illumination) and the delta ΣQ of the reversal learning cycle (with illumination), in GFP (filled green) and ARCH-expressing (white) mice. Delta ΣQ = ΣQ _end of the cycle_ - ΣQ _start of the cycle_. Gray lines represent the data from individual mice. Boxplots represent mean and interquartile range. Error bars are sem. **I)** Decision-values (ΔQ) variation during learning and reversal learning (with M2 bilateral illumination), separated by the black dashed line, in GFP- (full pink line) and ARCH-expressing mice (dashed pink line). Shaded areas, sem. **J)** Delta ΔQ of the learning cycle (without M2 illumination) and the delta ΔQ of the reversal learning cycle (with M2 illumination), in GFP (filled pink) and ARCH-expressing mice (white). Delta ΔQ = ΔQ_end of the cycle_ - ΔQ_start of the cycle_. Gray lines represent the data from individual mice. Boxplots represent mean and interquartile range. Error bars are sem.

This property of choosing an action based on the history of past events is at the heart of most RL models (Dayan & Daw, 2008; Katahira et al., 2015; Maia & Frank, 2011; Sutton & Barto, 2018; X. J. Wang, 2008). Thus, we designed a “*Q*-learning” RL model that captures accurately the animal’s lever-pressing strategy and reward rate (**Fig.3A**). Our model provided access to lever-pressing values *Q*, which represent the probabilistic expectation of obtaining a reward from an action. These values were updated by learning from the outcome (presence or absence of reward) according to Rescorla-Wagner-type equations (Ito & Doya, 2009) (**SupFig.3D**). The good prediction of the observed lever-pressing delays without fitting them (**SupFig.3E**), as well as the classical observation that the delay of action scales inversely with the sum of lever-pressing values (**SupFig.3F**), validate our modeling approach.

Here, we defined two task parameters ΔQ and ∑Q as the arithmetic combinations of the lever-pressing values over time (**Fig.3B**). ∑Q corresponds to the sum of the action values. As such, the change in ∑Q is independent of the order in which these action values are summed. In contrast, ΔQ is equivalent to DV and was defined here as the difference between the value of action 2 and the value of action 1. Consequently, ΔQ tends to equal -1 at the beginning of the learning cycle when the mouse presses its preferred but non-rewarding lever, and +1 when it successfully goes against its previous bias to maximize rewards (**Fig.3B**). By using a multiple regression analysis (Methods), we found that ΔQ and ∑Q influence task performance by controlling, respectively, the choice of action and the vigor at which that action will be initiated (**Fig.3C-F**), echoing previous results (Murakami et al., 2014; Tsutsui et al., 2016; A. Y. Wang et al., 2013). On the one hand, the observed delays between lever presentation and pressing were mainly negatively correlated with ∑Q (**Fig.3C**). The negative regression coefficients demonstrate that mice initiate lever-pressing faster when ∑Q is high (**Fig.3D**), which is consistent with motivation for rewards, regardless of the direction of choice. On the other hand, as expected from a DV, the fraction of success was positively correlated to the difference between the values of the reinforced and non-reinforced levers (the higher the value of one action in comparison to the other, the more likely it is to be chosen) (**Fig.3E, F**). In agreement, the impaired ability of ARCH mice to reverse lever-pressing choices when the activity of L2/3- PNs was inhibited during lever presentation (**Fig.1F, H**) was evidenced by a failure to update ΔQ in comparison to the GFP mice (learning; GFP: 1.22 ± 0.09; ARCH: 0.86 ± 0.16; n=11; p=0.38; reversal learning; GFP: -1.38 ± 0.16; ARCH: -0.68 ± 0.2; n=11; p=0.01; **Fig.3I, J**). In contrast, we found that GFP and ARCH mice had similar ∑Q without and with M2 illumination (learning; GFP: -0.33 ± 0.11; ARCH: -0.52 ± 0.16; n=11; p=0.63; reversal learning; GFP: 0.045 ± 0.04; ARCH: -0.09 ± 0.08; n=11; p=0.81; **Fig.3G, H**). These findings confirm that ∑Q is preferentially associated to a motivational aspect which sets the delay of actions (**Fig.3C, D**) and was not affected by M2 photo-inhibition (**Fig. 2A, B**). In contrast, ΔQ appears to be the main driver of reinforced choice and is directly updated by reward history.

### M2 guides actions by encoding behavioral task rule rather than upcoming choices

To understand how mice compute the DV ΔQ to solve the task, we used two-photon microscopy to image M2 neuronal activity through a glass cranial window (Aime et al., 2020) while mice learn and adapt their behavioral policy under the microscope (**Fig.4A**). We computed *ΔF/F_0_* from layer L2/3-PNs located in the right hemisphere and expressing the genetically encoded calcium indicator GCaMP6f (Aime et al., 2020; Chen et al., 2013). Individual PNs were automatically identified in images and tracked through multiple days (Methods). For the sake of consistency, the analysis was limited to ∼ 600 L2/3-PNs that were present in more than 80% of the learning and reversal sessions imaged (15.2 ± 6.6 sessions; 8.2 ± 3.9 learning sessions; 7 ± 3.4 reversal sessions; n=5 mice) (**SupFig.4A**).

**Figure 4:**
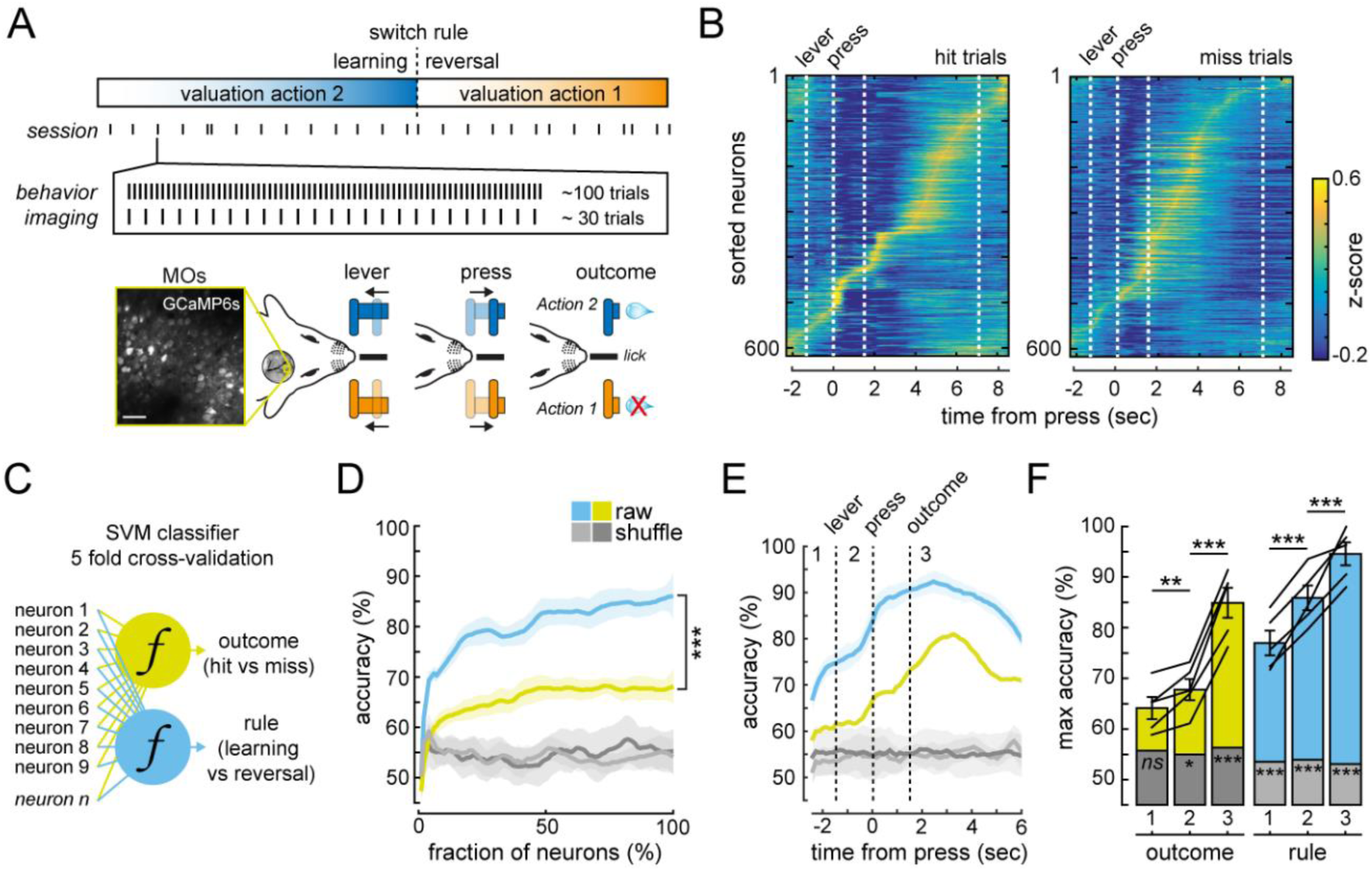
M2 neuronal activity decodes rule policy before lever-pressing. **A)** Schematic representation of the task policies and of the time windows of imaging of the neuronal activity. Each session consists on ∼100 trials, ∼30 of these were imaged. **B)** *Left panel*: heat map of the z-score of ΔF/F of GCaMP fluorescence for all averaged hit (rewarded, top panel) and miss (unrewarded, bottom panel) trials, from the neurons that pass the criteria (present in more than 80% of the imaged sessions). *Right panel*: similar to the left panel but for miss trials. **C)** Schematic representation of the two Support Vector Machines (SVM) classifiers used to infer the outcome and the task rule from the neuronal activity. **D)** Decoding accuracy (averaged across mice, +/- sem) as a function of the number of cells used for the SVM classifiers. **E)** Decoding accuracy (averaged across mice, +/- sem) of the SVM classifier for the outcome (green), the task rule (blue) and shuffled data (gray), throughout the trial. **F)** Max accuracy (averaged across mice, +/- sem) of the SVM classifier from the different periods of the trial (1, before lever presentation, 2, during lever presentation, 3, after outcome) for the outcome (green), the task rule (blue) and the shuffled data (gray). Error bars, sem.

First, we averaged *ΔF/F_0_* signals across all hit and miss trials from both learning and reversal learning sessions, and found that, individual L2/3-PNs displayed heterogeneous activity patterns with trial-averaged peak responses distributed over the entire trial period (**Fig.4B**). The difference in *ΔF/F_0_* signals between miss and hit trials within session is larger during the outcome period. This difference tends to be positively correlated with the increase in ΔQ that occurs between the beginning and end of each session (p=0.08) (**SupFig.4B**). This confirms that the activity of M2 L2/3-PNs is modulated by the reward and reflects the update of ΔQ. In contrast, the average z-score of *ΔF/F_0_* during the putative decision period (*i.e.* prior to lever-pressing) appeared similar in hit and miss trials (**SupFig.4B**). Nevertheless, L2/3-PNs in M2 could also operate in ensemble representations that potentially encode higher-level behavioral parameters in a more abstract manner (Rigotti et al., 2013). Thus, in order to better understand the neuronal representation of behavioral parameters during decision, we used a decoding approach to explore whether patterns of population activity before lever-pressing could serve action-selection by predicting the outcome (*i.e.* presence *vs* absence of reward) and/or implementing behavioral policy (*i.e.* learning *vs* reversal) (**Fig.4C-F**). To decode choice and policy representation from population activity (*ΔF/F_0_* from all cells without preselection), we constructed 2 support vector machine (SVM) classifiers which were applied to each time bin over the entire trial period (**Fig.4C**). We trained the classifiers on 80% of the trials (randomly chosen) regardless of the outcome but including the same fraction of the assigned trial types (*i.e.* hit *vs* miss - outcome; learning *vs* reversal - policy rule). We tested the classifiers on the remaining 20% of the dataset by comparing the classification results with actual trial types. This 5-time cross-validation procedure was repeated 1000 times with different training and test datasets, and the averaged prediction accuracy was computed across neurons (**Fig.4D**) and time (**Fig.4E, F**). To determine the chance level of the SVM classifiers, a permutation was done by randomly shuffling the assigned labels. Decoding accuracy improves as more neurons are added and reaches a maximum that is significantly higher for task rule policy than for the outcome (outcome: 68 ± 3.4 %; rule: 86 ± 4.5 %; n=5 mice; p<0.001; **Fig.4D**). The outcome and task rule policy, were both decoded from the entire neuronal population with significantly higher accuracy than chance after the lever was pressed and the outcome revealed (epoch 3; outcome raw: 84.918 ± 6.6737 %; outcome shuffle: 56.302 ± 11.807 %; p<0.001; rule raw: 94.574 ± 5.0592 %; rule shuffle: 53.108 ± 8.2808 %; p<0.001; n=5 mice) (**Fig.4D, E**). In contrast, during lever presentation (epoch 2) and inter-trial period (epoch 1), while the outcome was poorly or not at all decoded (outcome epoch 1 raw: 64.1 ± 4.8 %; shuffle: 55.7 ± 9.2; p=0.08; outcome epoch 2 raw: 68 ± 4.6 %; p=0.021; n=5 mice), task rule policy was decoded with high accuracy (rule epoch 1 raw: 77 ± 5.4 %; shuffle: 53.5 ± 6.7 %; p<0.001; rule epoch 2 raw: 86 ± 5.4 %; shuffle: 54 ± 5.8 %; p<0.001; n=5 mice). The accuracy of decoding before lever-pressing could not be explained by any licking-related activity (**SupFig.4C-F**), and is therefore unlikely to reflect any motor biases associated with licking. Hence, we propose that M2 L2/3-PNs govern which action to choose by preferentially encoding, before lever-pressing, the given behavioral policy rather than the upcoming decision.

### Neuronal representation of decision-value in M2 contributes to task policy

Since the behavioral policy probably results from a computational mixture of different task parameters, we generated a linear regression model to determine whether (and when within each trial) M2 neuronal responses encoded ΔQ, ∑Q, and/or the value of the chosen action, which were obtained from the RL model (**Fig.5A, B;** Methods). For each of these 3 regressors, the *p*-value associated with its β regression coefficient was used to identify the individual L2/3-PNs in M2 that were significantly modulated at different time bins across successive epochs of the task (**Fig.5A; SupFig.5**). In agreement with our decoding results (see **Fig.4**), the value of the chosen action was weakly encoded before the mice indicated their choice by pressing a lever and the outcome revealed, since the fraction of significantly modulated PNs oscillated around chance level during this period (chosen Q; raw: 6 - 13 % range; shuffle: 4 - 6 % range; n=5) (**Fig.5A**). This suggests that, in our task, the neural activity of M2 L2/3-PNs did not assign value to specific premotor activity, such as left or right direction. By contrast, a small but robust fraction of recorded L2/3-PNs were significantly modulated by ΔQ and ∑Q before lever-pressing (ΔQ: 12 - 42 % range; ∑Q: 11 - 34 % range; n=5) (**Fig.5A, B**). However, unlike ∑Q-coding neurons, identified ΔQ-coding neurons were predominantly positively regulated (β coefficients > 0; **SupFig.5D**), and their proportion was significantly higher than chance throughout the entire decision period (**Fig.5B**). In order to test whether the ΔQ signal in M2 plays a role in the encoding of task policy, four distinct SVM classifiers were constructed, each comprising a distinct set of cells whose activity was modulated by specific parameters from the reinforcement learning model (**Fig.5C**). In comparison to ∑Q-coding neurons, the activity of ΔQ-coding neurons before lever-pressing decoded task rule policy with a significantly higher level of accuracy (decoding accuracy relative to shuffled data; epoch 1; ΔQ: 1.8 ± 0.1; ∑Q: 1.6 ± 0.1; epoch 2; ΔQ: 1.85 ± 0.17; ∑Q: 1.66 ± 0.1; n=5; p=0.004; **Fig.5D**), despite the limited number of cells included in this population (**Fig.5E**). Our data indicate that, rather than other value-related signals, M2 L2/3-PNs preferentially signal ΔQ in the pre-press period, and this contributes to the encoding of task policy.

**Figure 5:**
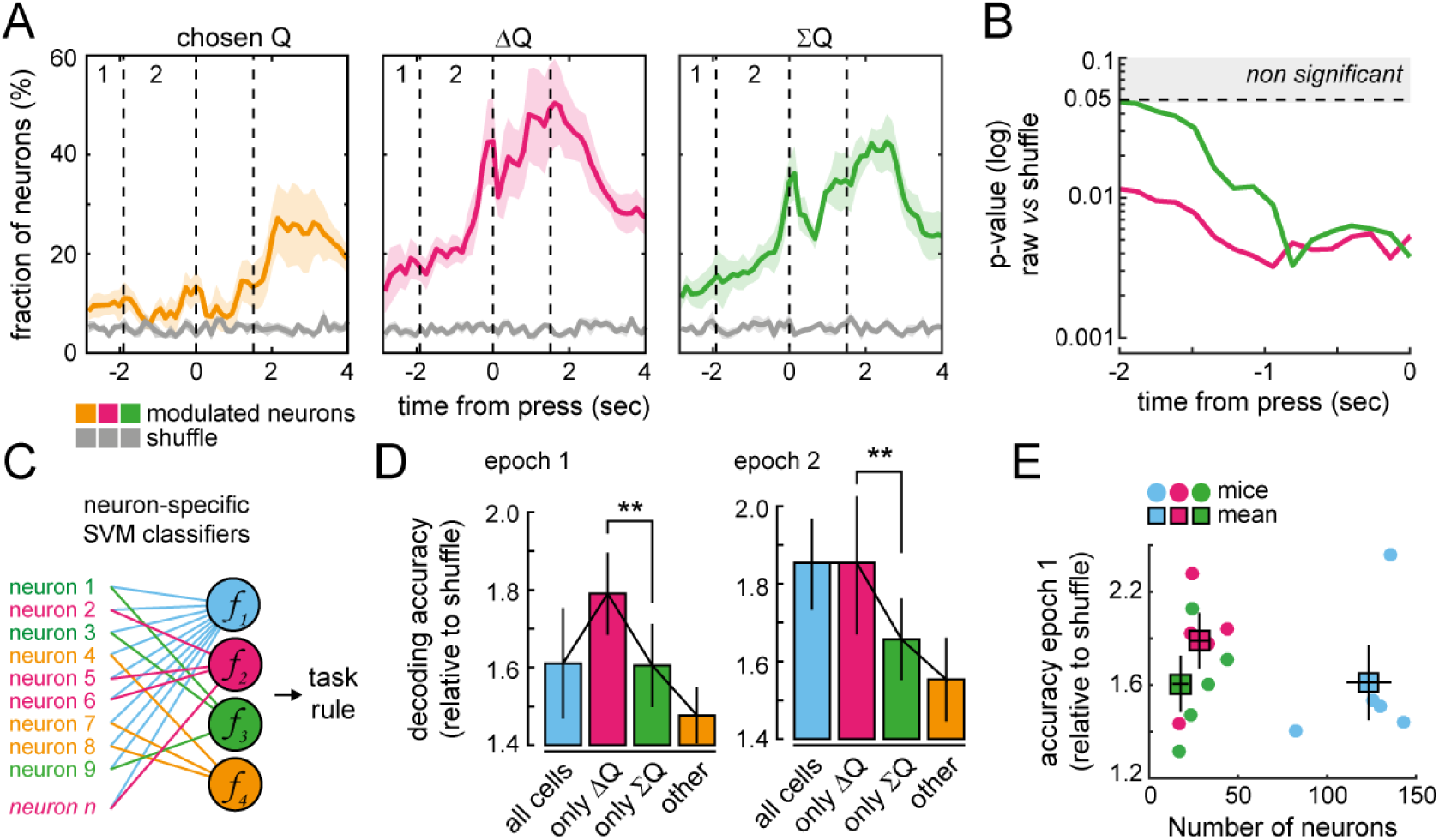
Differential contribution of value-modulated neurons to task rule representation. **A)** Fraction of neurons averaged across mice (+/- sem) modulated by the regressor *chosen Q* (orange), *ΔQ* (pink) and *ΣQ* (green) within each of the non-overlapping 150 ms time bins. Different periods of the trial are depicted (1, before lever presentation; 2, during lever presentation). Shuffled data in gray. **B)** p-value of the statistical comparison between the fraction of ΔQ (pink) and ΣQ (green) modulated neurons and shuffled data. **C)** Schematic representation of the Support Vector Machines (SVM) classifiers used to infer the task rule fromcell-specific modulated neuronal activity (blue, all neurons; pink, neurons modulated only by *ΔQ regressor*, green, neurons modulated only by ΣQ regressor; orange, neurons modulated by other than ΔQ and ∑Q regressors). **D)** Max decoding accuracy (relative to shuffled data, averaged across mice, +/- sem) of the SVM classifier from the different periods of the trial depicted in panel a, as a function of the identity of the modulated cells used in the classifier. Error bars, sem. **E)** Relation between the number and identity of cells included in the classifier (blue, all cells; pink, only ΔQ-cells; green, only ∑Q-cells) and the decoding accuracy during epoch 1 (before lever presentation). Each circle is a mouse, squares are the mean and error bars, sem.

### Neuronal activity related to decision-value in M2 is persistent and monotonically updated from trial to trial

In order for the brain to reliably select the action with the highest value using ΔQ, it must update this DV from trial to trial while maintaining it online within trials (Bari et al., 2019; Hattori et al., 2019). We tested these two properties, updating and stability, by quantifying how well the activity of M2 population before lever-pressing reflected DV (**Fig.6, SupFig.6**). ΔQ-coding neurons, which include all M2 L2/3-PNs with significant regression coefficients for ΔQ, gradually changed their firing rate during the decision period, which strongly correlates with ΔQ estimated from the model (r² = 0.8165; p<0.00001; **Fig.6A**). Notably, the neural activity of ΔQ-coding neurons increased monotonically during the learning phase, matching with the increase in ΔQ linked to the progressive selection of the rewarded action 2. Conversely, neural activity of the same neurons decreased with ΔQ during reversal while the value of the no longer rewarded action 2 decreased (**Fig.6B**). This is in contrast to the neural activity of ∑Q-coding neurons, which exhibited a lesser degree of correlation with changes in ∑Q (r² = 0.3730; p<0.00001; **Fig.6A, B**). As a consequence, the *t*-value of the regression weight was significantly higher for ΔQ than for ∑Q (**Fig.6C**), confirming the intimate relationship that exists during the decision period between DV and the activity of ΔQ-coding neurons across trials.

**Figure 6:**
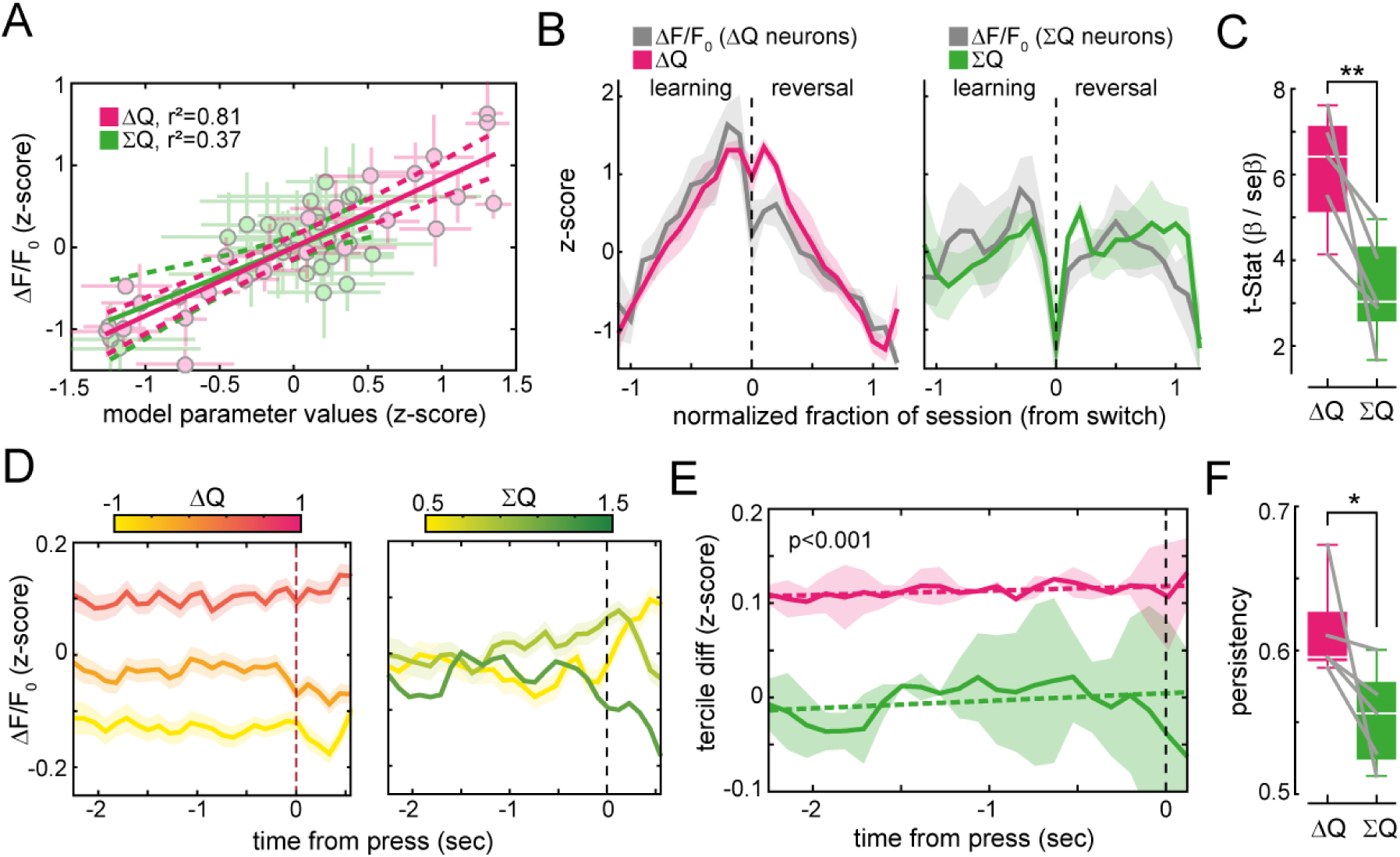
Monotic and persistent representation of decision variable in M2 neurons before lever-pressing. **A)** Correlation between ΔF/F_0_ (z-score) of ΔQ (pink) or ΣQ (green)-modulated neurons measured before lever-pressing and the values of ΔQ and ΣQ) obtained from the model. **B)** *Left panel*: z-score of ΔF/F (averaged across mice, +/- sem) of ΔQ-modulated neurons (grey) and of ΔQ (pink, averaged across mice, +/- sem) measured before lever-pressing across sessions, during learning and reversal, separated by the black dashed line. *Right panel*: z-score of ΔF/F of ΣQ-modulated neurons (grey) and of ΣQ (green, averaged across mice, +/- sem) across sessions, during learning and reversal, separated by the dashed black line. **C)** Statistic of the regression weight for ΔQ (pink) or ΣQ (green). Boxplots represent mean and interquartile range. **D)** ΔF/F (averaged z-score across mice, +/- sem) of ΔQ (*left panel*) and ΣQ (*right panel*) modulated neurons split by terciles of ΔQ and ΣQ, respectively. **E)** Mean difference between ΔF/F (z-score) across adjacent ΔQ (pink) and ΣQ (green) terciles, averaged across mice (+/- sem). Dotted lines, linear regressions. **F)** Persistency index reflecting the temporal stability of the neuronal activity of the neurons significantly modulated by ΔQ (pink) and ΣQ (green) during levers’ presentation. Boxplots represent mean and interquartile range.

We then quantified how stable the activity of ΔQ- and ∑Q-coding neurons was over the pre-press period within individual trials, which were classified according to model estimates (**Fig.6D-F**). First, we separated all trials collected from learning and reversal cycles into terciles of ΔQ or ∑Q, and averaged the neural activity of all ΔQ or ∑Q-coding neurons (**Fig.6D**). The difference between *ΔF/F_0_* averaged across adjacent terciles was significantly higher for ΔQ, showing that it discriminated neuronal activity across trials much better than ∑Q did (ΔQ: 0.11 ± 0.01; ∑Q: -0,004 ± 0.04; p<0.001, **Fig.6E**). Different ∑Q states are thus represented by similar pattern of neuronal activity. In addition, the persistency index, which describes formally the temporal stability of the activity of a given neuronal population as compared to the chance level (Hattori & Komiyama, 2022), is significantly higher for ΔQ- than ∑Q-coding neurons (ΔQ: 0.61 ± 0.01; ∑Q: 0.56 ± 0.02; n=5, p=0.04, **Fig.6F**). This analysis indicates that, in our task, M2 L2/3-PNs carry a signal related to DV, which remains persistent and more stable than all other model estimates throughout the pre-press period.

### The transition between decision-value states in M2 is linked to reversal performance

We then investigated whether the DV, that is ΔQ, specifically influenced behavioral flexibility during reversal learning. To do this, we restricted our analysis to post switch periods, during which the mice continued to exploit the wrong policy, explored the alternative option and eventually learned to exploit the correct new policy (period 1, 2, and 3, respectively; **Fig.7A**). Even if we confirmed that multi-collinearity did not affect the multiple regression analysis (**SupFig.7A**), linear models can still result in confounding mixtures of DV and other task parameters (Rigotti et al., 2013). To overcome this limitation, we used a dimensionality reduction technique that linearly compresses the firing rate of individual neurons into few task-dependent and task-independent dimensions (**Fig.7B**), while capturing the maximum amount of data variance (**SupFig.7B**) (Kobak et al., 2016; Passecker et al., 2019; Siniscalchi et al., 2016). Specifically, we used demixed principal component analysis (dPCA) (Kobak et al., 2016) to decompose the mix-selective population activity into: (1) the principal dimensions for ΔQ (ΔQ-axis), (2) trial type (*i.e.* miss and hit; choice-axis), (3) and the remaining independent dimension (free-axis) (**Fig.7B-D**). Because dPCA usually identifies dimensions for discrete variables (Kobak et al., 2016), the continuous variable ΔQ was split, averaged and categorized into three different ΔQ-states according to the reversal periods depicted in **Fig.7A**. This analysis revealed that the state of ΔQ is the main contributor of the variance of M2 neural activity as it explains more than 40 % of the total variability (ΔQ-axis: 46 ± 3 %; choice-axis: 13.7 ± 0.7 %; free-axis: 15 ± 4 %; n=5; **Fig.7C, D**). For all the mice analyzed, this first component with the largest possible variance always coincides with the ΔQ-axis (**SupFig.7C**). To determine the extent to which principal components segregate parameter dependencies, we shuffled trials by randomly permuting assigned labels (**Fig.7C, D**). The total variance explained by the independent dimension significantly increased as compared to raw data (free-axis; raw: 15 ± 4 %; shuffle: 45 ± 4 %; p<0.001; n=5). This comes at the expense of the ΔQ dimension, which explains now only around 20% of the variance (ΔQ-axis; raw: 46 ± 3 %; shuffle: 22.5 ± 1.6 %; p<0.001; n=5).

**Figure 7:**
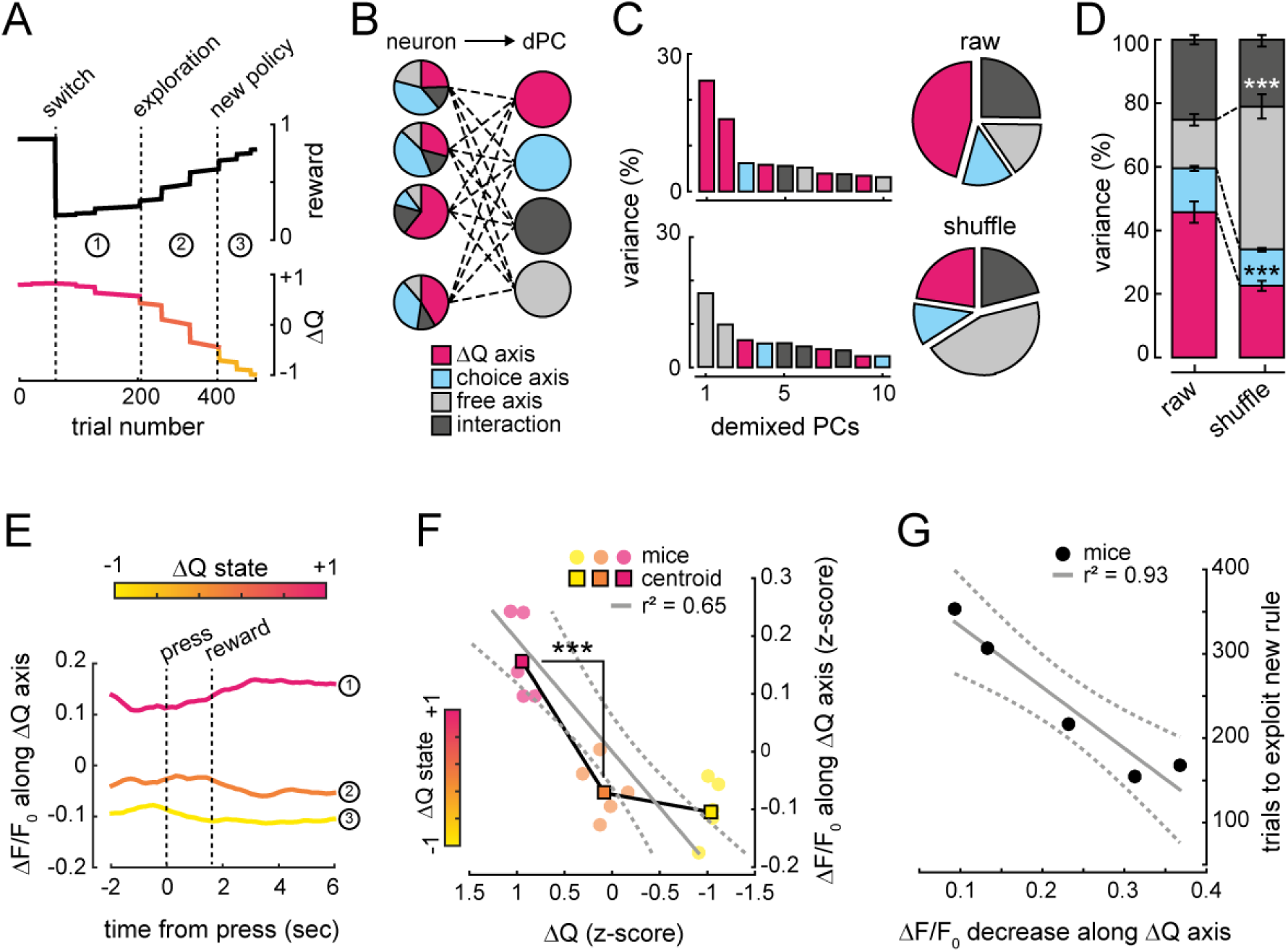
Reversal performance correlates with transitions of ΔQ states. **A)** Example of reward rate and ΔQ dynamic from one representative mouse. 3 different periods were used to average and categorize ΔQ into three different ΔQ-states during reversal: period 1 − the mice exploit the unrewarded lever, period 2 – the mice explored the alternative option and period 3 – the mice exploit the rewarded lever. **B)** schematic representation of the demixed principal component analysis (dPCA) for the unsupervised dimensionality reduction into demixed principal components (dPC). ΔQ-axis for ΔQ (pink), choice-axis for trial type (*i.e.* miss and hit; blue), free-axis for the remaining independent dimension (light grey)**. C)** Distribution of the variance in the first 10 PCs in task-dependent and task-independent dimensions, in the raw data (*top*) and the shuffled data (*bottom*). **D)** Percentage (mean +/- sem) of the variance in raw and shuffled data for the different axis as in b. **E)** The full data set from the same illustrative mouse in A is projected onto the dPCA decoder ΔQ axis, showing that the population activity is segregated along the first principal component (ΔQ axis) according to the 3 different ΔQ-states used for the dPCA and depicted in panel A. **F)** Correlation between ΔF/F_0_ of all recorded neurons during the decision period (*i.e.* before lever press) plotted along the ΔQ axis and the three ΔQ-states. **G)** Correlation between the total number of trials required to achieve expertise and the decrease of ΔF/F_0_ along the ΔQ axis and the two first ΔQ-states.

We next projected the neuronal activity onto the first principal component. This analysis reproduces the two main results obtained previously with methods based on linear regression (see **Fig.6**). First, *ΔF/F_0_* typically varies monotonically along the ΔQ-axis, indicating that neuronal activity decreases with decreasing ΔQ associated with different ΔQ-states during reversal (**Fig.7E**). Second, the firing pattern linked to each ΔQ-state remains persistent throughout the entire trial, particularly during the pre-press period (**Fig.7E**). We examined activity dynamics (trajectories) related to ΔQ-states by plotting the overall population activity within the neuronal manifold consisting of the ΔQ axis and the two task-independent axes with the greatest variance (**SupFig.7D**). It reveals that, despite the heterogeneity of single-neuron responses, the M2 population activity is orderly structured: trajectories are dominated by a circular geometry within each ΔQ-state, with each circle orthogonal to the ΔQ axis, and clearly segregated from the others along this axis (**SupFig.7D**). As a result, the trajectories never intersected, but probably translated from one to the other while ΔQ decreased during reversal, thereby allowing population activity to consistently reflect context (Russo et al., 2020) and update ΔQ necessary for the exploitation of the new rule (**SupFig.7E**). This hypothesis implies that the transition between ΔQ-state trajectories must be linked to the time needed to exploit the correct new policy. In agreement, the overall population activity computed from the decision period (*i.e.* before lever press) decreased significantly between the two first ΔQ-states (z-score *ΔF/F_0;_* state 1: 0.163 ± 0.03; state 2: -0.0650 ± 0.02; n=5; p < 0.001; **Fig.7F**). This transition drop is strongly negatively correlated with the total number of trials required to achieve expertise for each mouse (r² = 0.93, p=0.0077; **Fig.7G**). Therefore, mice that learnt the fastest showed the highest level of M2 activity along the ΔQ-axis, suggesting that the speed of the transition along the decision axis reflects the dynamics of the update of ΔQ during task performance.

## DISCUSSION

The secondary motor cortex (M2) in rodents is considered homologous to the supplementary motor area (SMA) in primates (Reep et al., 1987, 1990). The M2, like the SMA (Cadena-Valencia et al., 2018; Russo et al., 2020), supports highly integrated cognitive functions, including the inhibitory control of irrelevant action (Wardak et al., 2012), as well as the planning and initiation of voluntary action based on internal and/or external contextual factors (Barthas & Kwan, 2017; Ebbesen et al., 2018; Ebbesen & Brecht, 2017). As such, the elucidation of the neurocomputational mechanisms supported by the M2 is of paramount importance for the mitigation of repetitive and suboptimal behaviors that are frequently observed in various brain disorders (Everitt & Robbins, 2016; Izquierdo & Jentsch, 2012; Maia & Frank, 2011; Waltz & Gold, 2007). Our study expands on this previous research and provides new insights into the computational function of M2 in learning and reversal learning, especially when an abstract representation of the environment is required during learning due to the absence of clear sensory cues to predict rewarding actions. We have found that during the decision period, the population activity of M2 neurons accurately decodes the task rule. However, it does not discriminate between current choices, nor is it modulated by the value of the chosen action and the rate of licking. The lack of representation of these variables in M2 implies that these neurons are not specifically dedicated to a particular choice and are unlikely to contribute to any sensorimotor aspects of the task, at least prior to the execution of the action when decision is supposed to occur. In agreement, bilateral M2 photoinhibition does not affect the measured motor outputs, such as the delays and trajectories of the lever press and lick rate. These findings support the idea that M2 selects actions based on context rather than executing them, as previously suggested in rodents (Bari et al., 2019; Hattori et al., 2019; Hattori & Komiyama, 2022; Kawai et al., 2015; Siniscalchi et al., 2016) and primates (Russo et al., 2020; Tsutsui et al., 2016).

The lack of clear sensory cues before lever pressing suggests that other important internal factors are guiding action. Here, we found that head-fixed mice compute subjective values during reversal in a history-dependent manner. M2 neurons specifically and monotonically track the difference ΔQ across trials with well-structured neuronal trajectories unfolding along the ΔQ axis, thus unveiling a strong representation of ΔQ in M2. Consequently, the M2 neuronal population is well-suited to discriminate between different ΔQ states, thereby providing crucial insights into the task rule and the action that mice must perform in order to be rewarded. This highlights decision-value as an important internally driven contextual signal during reversal learning and confirms the critical role of premotor areas in encoding context in general, distinct from the function of pure motor areas (Hattori & Komiyama, 2022; Russo et al., 2018, 2020). During the pre-press period of each trial, persistent neuronal activity stably maintains ΔQ. This persistent pattern of neuronal activity may represent the ability to maintain ΔQ online and could therefore serve as a basis for short-term memory during the decision period. This could facilitate the conversion of graded, context-dependent changes in ΔQ into a binary choice, which is crucial for decision-making and motor planning (Brody et al., 2003; Murakami et al., 2014, 2017). Interestingly, the neural activity of ΔQ-coding neurons always increased monotonically with ΔQ during the learning phase as mice gradually abandoned the use of their preferred but non-rewarding forepaw (i.e., action 1). Because we defined ΔQ as the difference between the value of action 2 (non-preferred) and the value of action 1 (preferred) instead of the absolute difference between these values, it raises the possibility that the activity in M2 may be involved in suppressing irrelevant and impulsive actions or automatic behavioral habits. Such inhibitory control of executive function occurs in the SMA and pre-SMA of humans and primates (Wardak et al., 2012). Accordingly, the decrease in activity of ΔQ-coding neurons during reversal may reflect a disengagement of M2 as mice return to their initial bias.

We also found that M2 neurons are modulated by the sum ƩQ of action values, another decision variable which predicted lever press latencies with high accuracy. M2 bilateral photo-inhibition did not affect either, and the correlation is low between ƩQ and the neuronal activity of ƩQ-neurons during learning. As a consequence, the different states during learning and reversal are represented by similar patterns of activity of ƩQ-neurons. This suggests that the ƩQ signal in M2 is not necessary for the task, and that other brain circuits are involved in computing ƩQ (Turner & Desmurget, 2010; A. Y. Wang et al., 2013). This information can be then broadcasted during the decision period to other brain regions, including the M2 (Bari et al., 2019; J. H. Yang & Kwan, 2021). The function of the diffuse ƩQ copy signal remains unknown. However, it is consistent with the idea that ƩQ represents a general motivational factor that can enhance overall performance, rather than just modulating response initiation time (Turner & Desmurget, 2010; A. Y. Wang et al., 2013).

Decision variables have been observed previously in other tasks (Bari et al., 2019; Hattori et al., 2019; Hattori & Komiyama, 2022; Hirokawa et al., 2019; Tsutsui et al., 2016). Despite the computational similarities, they do not typically rely on M2, highlighting key differences between these tasks and ours. In most tasks, animals are passive and simply asked to lick either right or left depending on the presented stimulus. The difference in task-solving approach may be thus attributed to the use of two moving lever presses. This may have prompted the mice to solve the task in a more active way (Gilad et al., 2018), which was previously shown to depend on premotor areas such as the M2 or the SMA (Cazettes et al., 2023; Gilad et al., 2018; Russo et al., 2020; Sul et al., 2011; Tsutsui et al., 2016), in opposition to a passive behavior. In agreement, we found that bilateral M2 photoinhibition impairs reversal learning, which is consistent with previous findings that animals with M2 lesions were unable to reverse instrumental learned motor sequences (Gremel & Costa, 2013; Ostlund et al., 2009). Furthermore, it is possible that head-fixed mice solved the task in object space (Tsutsui et al., 2016), rather than in the action space observed with a single and static lick port.

On the other hand, our reversal task is also structurally different from the previous tasks, in which changes in reward probabilities occur within a single session, as these tasks typically result in rapid and accurate changes in choice allocation (Bari et al., 2019; Hattori et al., 2019; Hattori & Komiyama, 2022). In these studies, it took weeks of extensive training for the animals to optimally adapt their behavioral strategy to the structure of reward availability, suggesting that the mice had already learned the task rule, which was not the case here. In our task, head-fixed mice displayed a bias towards a particular side, leading to a persistence in exploiting the option with no reward during reversal learning. This indicates that, during learning, they do not immediately adopt an optimal win-stay/lose-shift behavioral policy, which classically uses the first previously unrewarded choice to rapidly explore another option. Instead, head-fixed mice integrate rewards from many previous trials. This information stored in memory is then used to learn how to make the best choice before self-initiating the lever press. This also suggests that the rule cannot be inferred from a single reward omission in the absence of an informative cue, thus increasing the difficulty of the task even with deterministic reward contingencies. However, actions remained deliberate and goal-directed, as reward devaluation drastically reduced action rate (Bouton & Balleine, 2019).

What mechanisms and circuits convert the decision value into a binary motor command remains an open and critical question. It is possible that this arises through long-range loops between multiple brain regions, similar to memory-guided licking tasks in mice. Indeed, previous research in rodents showed that M2 neurons link instructive sensory information with movement execution, even when the action is delayed after the sensory input has ended (Brody et al., 2003; Erlich et al., 2011; Guo et al., 2014; Komiyama et al., 2010; Kopec et al., 2015). During the delay, which is comparable to the decision period in our task, the neuronal activity in M2 gradually changes and predicts particular future movements (Inagaki, Chen, Daie, et al., 2022). In these tasks, planning and executing movement involves multiregional neural circuits including the thalamus, midbrain, cerebellum, and basal ganglia (Inagaki, Chen, Ridder, et al., 2022; Svoboda & Li, 2018). On the other hand, because we recorded superficial L2/3 neurons we do not rule out the possibility that conversion occurs locally with the help of L5 neurons projecting to downstream motor structures (Economo et al., 2018; Murakami et al., 2017). This would require the coexistence across M2 layers of multiple attractor networks with different architecture (Miller & Katz, 2010; Murakami et al., 2014, 2017; Svoboda & Li, 2018). To address all of these possibilities, it is now necessary to combine large-scale recordings of input/output-specific neuronal populations with transient optogenetic perturbations. This will allow us to assess the contribution of each possible network at play during behavior (Svoboda & Li, 2018).

## METHODS

### Contact for Reagent and Resource Sharing

Correspondence and reasonable requests for materials and data supporting this study should be addressed to (Lead contact)

### Experimental Model and Subject Details

A total of 27 adult males C57BL/6J from Charles River, 2-7 months old, were used in this study (5 for calcium imaging and 22 for optogenetics). Mice were housed with littermates (3–4 mice per cage) in ventilated cages under a normal 12 h light/12 h dark cycle, temperature was maintained between 19 °C and 23 °C and humidity between 50% and 65%. The mice were housed in enriched cages and provided with food and water ad libitum. During the head-fixation phase (training, recordings and optogenetics), the mice were water-restricted (beginning approximately 10 days after surgery), with water available only during the task or afterwards until a total of 1 mL per day. On days when no training or recording occurred, the mice were given 1 mL of water. All experimental procedures were approved and performed in accordance with the local ethics and welfare Committees guidelines (N°50DIR_15-A) and by the French Ministry for Agriculture/Research (APAFIS #21728; APAFIS #18907; APAFIS #38205), in agreement with the European Communities Council Directive of September 22th 2010 (2010/63/EU, 74).

### Method Details

#### Surgery for calcium imaging and optogenetic

All surgeries used standard aseptic procedures. Mice were deeply anesthetized with 4% of isoflurane (by volume in O_2_) for 5 min, further anesthetized with an intraperitoneal injection of a mix containing medetomidine (sededorm, 0.27 mg.kg^-1^) and buprenorphine (0.05 mg.kg^-1^) in sterile NaCl 0.9%, and mounted in a stereotaxic apparatus (RWD). Mice were kept on a heating pad and their eyes were covered with eye ointment ophthalmic gel. The scalp was shaved and disinfected with 70% ethanol and betadine. Local analgesia was achieved by application of 100 ml of lidocaïne (lurocaïne, 1%) injected subcutaneously in the scalp. 40 μL of dexamethasone (dexadreson, non-steroidal anti-inflammatory and analgesic drug, 5 mg.kg^−1^) was administrated intramuscularly in the quadriceps to prevent inflammation. The top skin (a circumference with around 1 cm of diameter) was removed from the dorsal skull with scissors and the skull was cleaned and dried with sterile cotton swabs. For stereotaxic injections the bregma and lambda were aligned (x and z). The injections targeted the layer 2/3 of the M2 (three injection sites: between +2.3 mm and +3 mm AP; between -0.2 mm and -0.3 mm DV; ± 1.5 mm ML from bregma). A total of 300 nL (100 nL per site) of virus was injected at a maximum rate of 60 nL.min^−1^, using a glass pipette (Wiretrol, Drummond) attached to an oil hydraulic manipulator (MO-10, Narishige). AAV1.CaMKII.GCaMP6f.WPRE.SV40 (Addgene, USA) was injected for calcium imaging after craniotomy, and for optogenetics AAV9.CAG.ArchT.GFP.WPRE.SV40 for ARCH-expressing mice or AAV9.CamKII0.4.eGFP.WPRE.rBG in GFP-expressing mice (Addgene, USA) was injected. After injections, the viruses were allowed to diffuse for at least 10 min before the pipette was withdrawn. Mice were then either prepared for cranial window or optical fiber implantation. The bone was scraped with a delicate bone scraper tool and covered with a thin layer of dental cement (Jet Repair Acrylic, Lang Dental Manufacturing).

The cranial windows were implanted as previously described (Aime et al., 2020; Gambino et al., 2014). Briefly, after skull’s exposure a ∼ 5 mm plastic chamber was attached on the area of interest and a 3 mm craniotomy was drilled on the right hemisphere above M2 (around +3 and +1 mm AP; 3+ and 1+ mm ML), with a pneumatic dental drill (BienAir Medical Technologies, AP-S001) leaving the dura intact. After viral injection, the craniotomy was filled with sterile saline (0.9% NaCl) and sealed with a 3 mm glass cover previously sterilized and soaked in 0.9% NaCl sterile solution. The chamber and the cover slip were bonded to the skull using acrylic glue and dental cement (Jet Repair Acrylic, Lang Dental Manufacturing). For optical fibers implantation, two small craniotomies were drilled above M2 (around +2.8 mm AP; ±1.5 mm ML) and the optical fibers (CFML22U-20, Thorlabs) were implanted above the craniotomies previously soaked in 90% ethanol and afterwards in 0.9% NaCl sterile solution. The optic fiber with guided cannula for optogenetics (CFML22U, Thorlabs) were cleaved with a fiber optic scribe (S90R, Thorlabs) at 0.2 mm and implanted bilaterally inside the two small craniotomies performed on top of M2 (+2.8 mm AP; ±1.5 mm ML from bregma) and barely touched the dura (as to avoid damaging superficial cortical layers). The optical fibers were stabilized with glue and dental cement.

At the end of the surgery, anesthesia was reversed by a sub-cutaneous injection of a mixture containing atipamezole (revertor, 2.5 mg.kg^-1^) and buprenorphine (buprecare, 0.1 mg.kg^-1^) in sterile 0.9% NaCl. 7-10 days post-surgery a custom-made stainless-steel head stage (stainless steel, 2 × 0.5 cm, 1g) was positioned on top of the dental cement above the cerebellum and dental cement (Jet Repair Acrylic, Lang Dental Manufacturing) was added to fix the head bar in position.

#### Behavioral apparatus and training

The mice were placed on a behavioral apparatus custom-designed by the company Imetronic. It consists of a linear metal grid with metal walls on the sides, two metal pads (right and left) in the frontal part of the grid, a lick port connected to a pump that controls the water flow to the lick port, and two retractable metal levers (right and left) below the lick port and in front of the metal grid, where the head-fixed mouse stands. The software that controlled the apparatus was also responsible for detecting in real-time forepaw movements in the pads, licks, and lever presses (POLY, by Imetronic). The apparatus was placed within a soundproof black box, which was monitored throughout each session by an IR camera (Sony IR color CCD camera, horizontal resolution 700tvl, lens 8:2.5 mm) positioned on the apparatus’s side.

Mice were handled by the experimenter for at least 7 days, starting from 7 days post head bar implantation, during the water restriction and before the first habituation session. At the beginning of the habituation, mice were acclimatized to the head fixation and to the lick-port by receiving water reward drops (10 μL) available at the lick-port at a frequency of 0.09 Hz. Mice were head restrained until they stopped licking for reward or until they consumed a total of 1 ml of water. After mice learned to lick for water reward (typically after one or two sessions), the training task consisted of an easier version of the task, during which a mix of two protocols were used: 1) both levers were presented simultaneously and both triggered a water reward 1.5 sec after right or left press, 2) right and left levers were presented alternatively and both triggered a water reward 1.5 sec after right or left press. That way, mice learn to press the levers and the association between pressing the lever and water reward. Performance improvement was indicated by an increase in the number of trials in 1 hour of training (max 100 trials), which was dictated by a decrease in the average time to press the levers after presentation within a trial. Mice were trained for several (between 3 and 14) consecutive days on the training task until they reached 100 trials in 1 hour of training before the learning sessions begin.

#### Learning – reversal learning task design

In the bandit task, two retractable levers (right and left) could be reached and pressed with the right or left forepaws respectively, when presented. Press consists of a 100 msec continuous press of a lever (∼3.3 g) and after press the levers were immediately retracted. Mice could lick the lick-port at any given period of the trial, but water rewards were only available at 1.5 sec after lever press. After water reward delivery there was an inter-trial interval of 7 sec, before the next trial began with both levers presented again. There were no explicit cues that allowed discriminating when the water was being delivered, neither when the levers were presented. During learning, both levers were simultaneously presented but only the press of one lever (the one to which each mouse had less bias) was triggering a water reward. During reversal the lever whose press triggered a reward was the opposite of learning. Mice were trained for several (between 3 and 14) consecutive days on the learning task before the reversal sessions begin. Expertise was defined as 3 consecutive days with an average of 75% of rewarded choices.

The reward probability of a hit trial (rewarded) was 100% and of a miss trial (unrewarded) was 0%. The reason for choosing these statistics is that they correspond to a low level of environmental uncertainty, which allows the mice to learn the task faster and to remain highly motivated during the sessions, thus yielding a large number of behavioral trials per session. After expertise *ad libitum* water was provided in the home cage for 48 hours and the frequency of the lever press was assessed with a session of training task 1.

#### 2-photon calcium imaging

Head-fixed awake mice were placed and trained under the microscope every day for around 4 days, until they drank 1mL of water under head-fixation, before training started. Imaging started 21 to 35 days after virus injection, using an in vivo non-descanned FemtoSmart 2PLSM (Femtonics, Budapest, Hungary) equipped with a x16 objective (0.8 NA, MRP07220, Nikon). The MES Software (MES v.4.6; Femtonics, Budapest, Hungary) was used to control the microscope, the acquisition parameters, and the TTL-driven synchronization between the acquisition and behavior apparatus. The GCaMPs were excited using a Ti:sapphire laser operating at 910 nm (Mai Tai DeepSee, Spectra-Physics) with an average excitation power at the focal point lower than 50 mW. Time-series images were acquired at ∼37 Hz. We imaged on average 11000 frames (∼300 sec) during each session, and no visible photo-bleaching was observed. Each imaging session was acquired for a random 5 min of each behavioral session.

#### Optogenetic stimulation

To optically stimulate ARCH-expressing neurons, we used a 561 nm light from a diode-pump solid-state (DPSS) laser (SDL-LH-1500, Shanghai Dream Lasers Technology). Light was emitted from the laser through an optical fiber patch cable (M83L01, Thorlabs) and connected through a ceramic split mating sleeve (ADAL1, Thorlabs), to the chronically implanted optic fiber cannulas. The power of the laser was calibrated using an optical power meter kit (PM200, Thorlabs). During the task, the optical stimulation (10 mW) was turned on during each lever presentation of the reversal cycle. Light delivery was triggered by a pulse-stimulator (Master-9, A.M.P.I), able to synchronize laser with lever presentation and retraction, light delivery started when the levers were presented and stopped when the mouse pressed any lever for 100 msec inducing lever retraction.

#### Histology and fiber localization

To evaluate the viral expression profiles and optical fiber location in M2, after behavior completion, mice were deeply anesthetized with pentobarbital (300 mg.Kg^-1^) and perfused with 4% paraformaldehyde. The brain was extracted and fixed for 24 h in paraformaldehyde at 4 °C, and then washed with phosphate-buffered saline. The brain was sectioned at 50 µm, mounted on glass slides and stained with DAPI. Images were taken at ×20 magnifications for each section using a Hamamatsu Nanozoomer at two different excitation wavelengths (405 nm and 488 nm for DAPI and GFP, respectively). To determine the approximate location of the optical fibers implantation and GCaMP6s, ARCH and GFP expression, we added a mask from the mouse brain atlas (Franklin and Paxinos) on top of our imaging of brain slices. A visual guess was made to find the coordinates from the Paxino Mouse Brain Atlas by comparing structural aspects of the histological slice with features in the atlas, for each mouse for the slice with the highest expression of the virus.

### Quantification and Statistical Analysis

#### Preprocessing of imaging and behavioral data

Fluorescence images were preprocessed using Suite-2P (Pachitariu et al., 2016) in Matlab (Mathworks). Suite-2P corrected XY motion artifacts over time using an elastic registration, segmented the somas of individual neurons based on their GCaMP signal, and extracted the fluorescence signal of each soma per frame. Cell segmentation was manually curated using the GUI of Suite-2P and excluded ROI were not further analyzed. Registration of neurons between session was done manually, and we only selected for analysis neurons that were present in more than 80 % of all the imaged sessions. Each session consisted of about 100 behavioral trials, 30 of which were imaged. The behavioral and imaging data were synchronized through the transmission of 6 TTL signals sent to the microscope. Each signal was digitalized at 1 kHz and stored as metadata, corresponding to a specific behavioral parameter: two for each right and left moving lever, two for the press of each lever, one for the licks detected at the lick-port, and one for the pump controlling the drop of water reward.

The ΔF/F_0_ was calculated for each neuron as described previously (Aime et al., 2020) and normalized (z-score) throughout the entire session, independently of one another, before being split into trials. We defined each trial as the period between lever presentation -5 sec and water delivery + 5 sec. As the task was self-initiated, the interval between lever presentation and lever pressing varied between trials, with each trial exhibiting a distinct duration. In order to align events across trials for each mouse, the trials were re-stretched by linearly time-warping the signal acquired during lever presentation and lever-pressing. This was done to align the time axis with the median interval between all lever presentations and lever-presses, for each mouse independently. Only trials that were shorter than two seconds were time warped and further used for post-processing analysis to ensure that time warping did not introduce any artifacts.

#### Tracking of behavior

The behavior (choice, press delay and lick frequency) was analyzed according to the time points of each event provided by POLY (Imetronic software). Videos of the animal performing the task were acquired using an infra-red camera (Sony IR color CCD camera, horizontal resolution 700 TVL, Len 8:2.5 mm). For forepaw trajectory analysis, the movements of (1) the forepaws during lever press, (2) the levers during presentation and retraction, were extracted and tracked from videos using DeepLabCut (Mathis et al., 2018). A total of 663 frames with lever presentation and retraction were selected and labeled (2 points to track one lever, 1 point to track the other lever and 1 point to track each forepaw) to train the network.

#### History-based and reinforcement learning models

In order to predict current correct choice behavior and quantify the extent to which historical factors influence the reward rate, we calculated logistic regression according to:

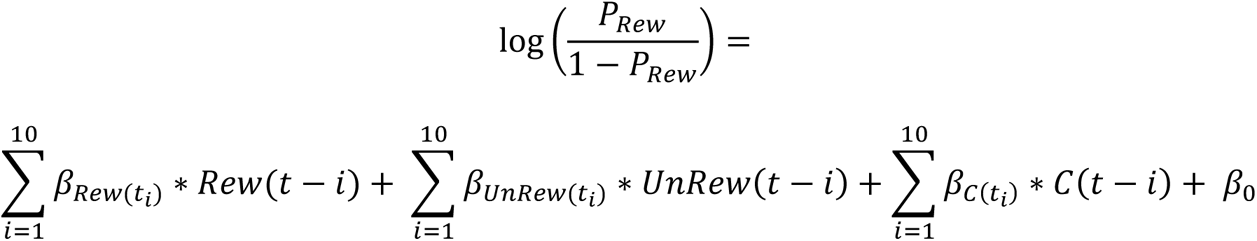

We used 3 predictor types: *Rew*(*t*−*i*) is the rewarded choice history on trial *t*−*i* (+1 if rewarded lever 2 choice, −1 if rewarded lever 1 choice, 0 otherwise), *UnRew*(*t*−*i*) is the unrewarded choice history on trial *t*−*i* (+1 if unrewarded lever 2 choice, −1 if unrewarded lever 1 choice, 0 otherwise), *C*(*t*−*i*) is the outcome-independent choice history on trial *t*−*i* (+1 if lever 2 choice, −1 if lever 1 choice). *`β`* represents the regression weight of each history predictor, and *`β`*_0_ is the history-independent constant bias term. The accuracy of prediction was performed by a 2-fold cross-validation and compared to the result of a logistic regression using a lever bias term. For reinforcement learning, we used a standard Q-learning model to describe the lever-pressing strategy of animals through learning: the lever-pressing value (*Q*_1_(*t*) and *Q*_2_(*t*) for lever 1 and 2, respectively) reflected the probabilistic expectation of obtaining a reward when taking the action. Accordingly, they were updated when actions were taken and their outcome evaluated according to Rescorla-Wagner-type equations (Rescorla and Wagner, 1972). In particular, if L1 is pressed:

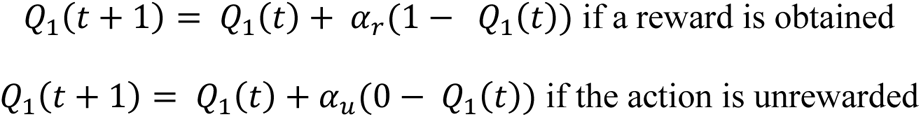

By setting different update parameter values for successes and failures (*α*_*r*_ ≠ *α*_*u*_) the model allows perseveration to take place (Ito & Doya, 2009). Moreover, when L1 is not pressed, we apply a forgetting update *α*_*f*_ (Ito & Doya, 2009):

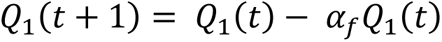

Similar equations apply to L2. Although pressing together L1 and L2 (double-pressing) occurred transiently through learning, it never led to reward and, as a result, should never be reinforced. Consequently, it discarded a three-state model where double-pressing is an independent action which can be selected. Instead, double-pressing was considered to be the simultaneous selection of pressing both L1 and L2.

In reinforcement learning models, probabilistic action selection is typically computed using a function of the ensemble of action values (Qs). We postulate that action selection is based on a race model between actions, wherein the delay for taking an individual action is linked to its value as previously described (Turner & Desmurget, 2010; A. Y. Wang et al., 2013). We describe the probabilistic pressing of lever L1 as the probability that its delay of action (*T*_1_) is shorter by a margin larger than *δt* than the delay for pressing L2 (*T*_2_):

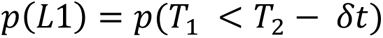

We chose to describe delays of action as random variables following an exponential distribution:

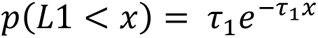

The inverse mean (*τ*_1_) is computed from the scaled lever value: *τ*_1_ = *a*_1_ + *bQ*_1_, with *a*_1_a bias factor modeling the intrinsic preference of animals for the lever (smaller is better), and *b* a scaling factor from lever value to delay. Using this model, we computed the lever pressing probabilities as:

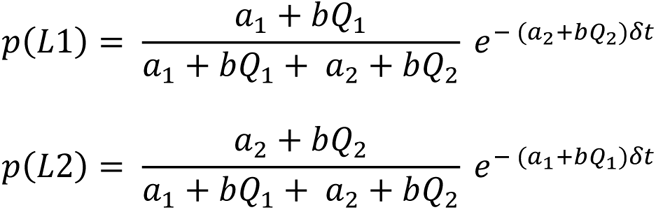

From the actions value Q_1_ and Q_2_, we defined the task parameters ΔQ and ∑Q as the arithmetic combinations of the lever-pressing values over time: ∑Q corresponds to the sum of the action values, and ΔQ is the difference between the value of action 2 and the value of action 1

#### Validity of reinforcement learning model

It has been observed previously that the delay for action is correlated with the total value of actions ∑Q, with larger ∑Q resulting in shorter delays. In our model, the predicted overall delay of action does not depend on the identity of the action taken and has a mean of 1⁄(*a*_1_ + *a*_2_ + *b*(*Q*_1_ + *Q*_2_)), scaling thus with ∑Q. Therefore, in order to assess the validity of the reinforcement learning model, as well as the influence of ΔQ and ∑Q on behavioral parameters of the current choice, we used the following multiple linear regression models:

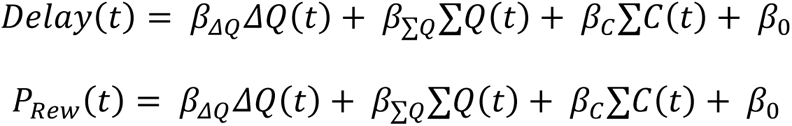

Where *Delay(t)* is the delay between lever presentation and lever retraction, *P_Rew_*(t) is the probability of reward, and *C*(*t*) is the outcome-independent choice on trial *t* (+1 if pressing lever 2, −1 if pressing lever 1). The regression coefficients β for the 3 regressors ΔQ, ∑Q and C were estimated simultaneously.

#### Multiple linear regression analysis of neuronal activity

We performed multi-regression analysis to quantify the fraction of neurons that were significantly modulated by ΔQ (ΔQ-coding neurons) and ∑Q (∑Q-coding neurons). We first averaged the *ΔF/F_0_* signal of each neuron with non-overlapping moving windows (bins of 250 msec), and then used for each time bin the following regression model:

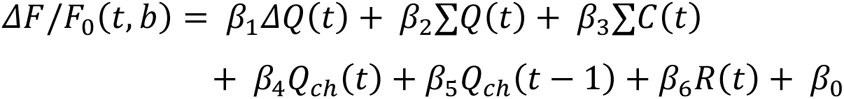

Where *ΔF/F_0_(t, b)* corresponds to the averaged *ΔF/F_0_* at time bin b for each neuron, *ΔQ(t)* is the difference of value between lever 2 and lever 1 on trial *t*, *∑Q(t)* is the sum of value on trial *t*, *C*(*t*) is the outcome-independent choice on trial *t* (+1 if pressing lever 2, −1 if pressing lever 1), *Q_ch_(t)* is the value of the chosen action on trial *t*, *Q_ch_(t-1)* is the value of the chosen action on previous trial *t-1*, *R(t)* is the reward outcome on trial *t* (+1 if reward, −1 if no reward). β_1…6_ are the corresponding regression coefficients, which were tested for statistical significance with two-sided t-test. Since the regression coefficients are estimated simultaneously, we ensured that multicollinearity between the regressors did not affect the results by applying the following constraints. 1) First, we calculated for each neuron and each time bin the variance inflation factor (VIF), a standard measure for multicollinearity. This was done to ensure that the different independent regressors in the multiple regression model were not correlated. The mean VIF for *β_1_, β_2_, β_3_, β_4_, β_5_, β_6_* were 1,26 ± 0.07, 2.84 ± 0.5, 1.94 ± 0.15, 2.91 ± 0.7, 2.21 ± 0.45, and 1.94 ± 0.15, respectively (n=5 mice). 2) In order to further minimize the effects of multicollinearity, only those regression coefficients that were statistically significant were used. 3) Finally, we compared these statistically significant regression coefficients with those obtained by permuting the assigned labels (shuffled data). Only cells with raw regression coefficients that were statistically different (two-tailed t-test) from the shuffled data were considered modulated by any given regressor.

#### Decoding of behavior from population activity

We used SVM classifiers to decode binary choice (hit *vs.* miss trials) and binary policy representation (learning *vs.* reversal learning) from neuronal population activity. This was done for each mouse independently, using all trials acquired during all behavioral sessions. However, we homogenized the number of trials so that they were equivalent for each grouping category. For example, if the total number of hits exceeded the number of misses, we randomly selected an equal number of hit trials to build the binary choice classifier. To train the classifier, we generated, for each time point of a trial, a *t* x *n* “training” matrix containing the *ΔF/F_0_* activity of *n* neurons for each *t* trials. This “training” matrix contained 80% of all trials which were randomly selected. We generated another “label” matrix that contained the corresponding binary labels associated to each trial (which were later randomly permutated to create shuffled data). This step was repeated 1000 times to generate 1000 different training matrices with different randomly selected trials. To test the effect of neuronal population size, *n* was increased from 1 to the maximum number of neurons recorded in each mouse. The SVM model was built using the ‘fitcsvm’ built-in function implemented in Matlab (Version R2014b, Mathworks), and trained by the predictor “training” and class labels “label”. We used a linear kernel and the default optimization parameters for finding the SVM separating hyperplane. This SVM model was then used to classify the remaining 20% of trials using the Matlab built-in ‘predict’ function. We quantified decoding accuracy as the percentage of correctly classified trials, averaged over all 1000 consecutive iterations.

#### Demixed principal component analysis (dPCA)

We used dPCA as previously described (Kobak et al., 2016; Passecker et al., 2019; Siniscalchi et al., 2016) to decompose the activity of the M2 neuronal population into a few demixed components that capture most of the variance while keeping task parameters, such as decision states and choices, as separated as possible. To do this, we generated, for each mouse, a single multidimensional *n* x *S* x *C* x *T* x *t* matrix containing the average *ΔF/F_0_* activity for each neuron *n*, each ΔQ state *S*, each time point *T* of the trial *t*. We defined *C* as hit and miss trials, and *S* as the average ΔQ for quantiles of the ΔQ values for the probability [0, 0.33, 0.66, 1]. All trials shorter than 2 seconds, which were previously time-warped to make sure they were the same length, were concatenated. This input matrix was then analyzed and demixed by using the MATLAB scripts provided by Kobak et al., 2016 (https://github.com/machenslab/dPCA).

## Acknowledgements and Funding

We thank H. El Oussini, P. Costet and C. Martin (PIV-EOPS & PIV-EXPE, Univ. Bordeaux, CNRS) for support with animal husbandry; S. Valerio (AquiNeuro) and the Bordeaux Imaging Center (BIC, Univ. Bordeaux, CNRS, INSERM, BIC, US4, UAR 3420) for their technical expertise and support, and all the members of the Gambino laboratory for technical assistance and helpful discussions. We thank the PIV-EOPS and PIV-EXPE facilities of the IINS. We thank K. Deisseroth and Stanford University, E. Boyden and MIT, L.L. Looger and D. Kim from the GENIE project, and K. Svoboda at the Janelia Farm Research Campus (HHMI) for distributing viral vectors. During the preparation of this work the authors used with caution “DeepL Write” only in the writing process in order to improve language and readability. After using this tool, the authors reviewed and edited the content as needed and take full responsibility for the content of the publication.

This project has received funding from (to FG): the European Research Council (ERC) under the European Union’s HORIZON research and innovation program (ERC-CoG-2021; MOTORHEAD; grant agreement 101043602); the French National Research Agency (grant agreement ANR-21-CE37-0010-SyTUNE; grant agreement ANR-22-CE37-0015-THALACOR; ANR-22-ERCC-0010-MOTORHEAD); the University of Bordeaux; and the Region Nouvelle Aquitaine. EA is supported by a Marie Skłodowska-Curie individual fellowship (HORIZON-MSCA-2023-PF-01) under the European Union’s HORIZON research and innovation program (DANAL; grant agreement 101155516). NC was supported by a Marie Skłodowska-Curie individual fellowship under the European Union’s Horizon 2020 (H2020-MSCA-IF-2017) research and innovation program (AXO-MATH; grant agreement 798326). This work was supported by the Bordeaux Neurocampus core facilities (LabEx BRAIN; grant ANR-10-LABX-43). The Bordeaux Imaging Center was supported by the French National Research Agency (grant agreement ANR-10-INBS-04).

## REFERENCES

Aime, M., Augusto, E., Kouskoff, V., Campelo, T., Martin, C., Humeau, Y., Chenouard, N., & Gambino, F. (2020). The integration of Gaussian noise by long-range amygdala inputs in frontal circuit promotes fear learning in mice. ELife, 9. 10.7554/elife.62594

Balewski, Z. Z., Knudsen, E. B., & Wallis, J. D. (2022). Fast and slow contributions to decision-making in corticostriatal circuits. Neuron, 110(13), 2170–2182.e4. 10.1016/J.NEURON.2022.04.005

Banerjee, A., Parente, G., Teutsch, J., Lewis, C., Voigt, F. F., & Helmchen, F. (2020). Value-guided remapping of sensory cortex by lateral orbitofrontal cortex. Nature, 585(7824), 245–250. 10.1038/s41586-020-2704-z

Bari, B. A., Grossman, C. D., Lubin, E. E., Rajagopalan, A. E., Cressy, J. I., & Cohen, J. Y. (2019). Stable Representations of Decision Variables for Flexible Behavior. Neuron, 103(5), 922–933.e7. 10.1016/j.neuron.2019.06.001

Barthas, F., & Kwan, A. C. (2017). Secondary Motor Cortex: Where “Sensory” Meets “Motor” in the Rodent Frontal Cortex. Trends in Neurosciences, 40(3), 181–193. 10.1016/j.tins.2016.11.006

Bissonette, G. B., Martins, G. J., Franz, T. M., Harper, E. S., Schoenbaum, G., & Powell, E. M. (2008). Double dissociation of the effects of medial and orbital prefrontal cortical lesions on attentional and affective shifts in mice. The Journal of Neuroscience: The Official Journal of the Society for Neuroscience, 28(44), 11124–11130. 10.1523/JNEUROSCI.2820-08.2008

Bohn, I., Giertler, C., & Hauber, W. (2003). Orbital prefrontal cortex and guidance of instrumental behaviour in rats under reversal conditions. Behavioural Brain Research, 143(1), 49–56. 10.1016/S0166-4328(03)00008-1

Bouton, M. E., & Balleine, B. W. (2019). Prediction and control of operant behavior: What you see is not all there is. Behavior Analysis (Washington, D.C.), 19(2), 202–212. 10.1037/BAR0000108

Brody, C. D., Romo, R., & Kepecs, A. (2003). Basic mechanisms for graded persistent activity: Discrete attractors, continuous attractors, and dynamic representations. Current Opinion in Neurobiology, 13(2), 204–211. 10.1016/S0959-4388(03)00050-3

Brown, V. J., & Tait, D. S. (2014). Behavioral Flexibility: Attentional Shifting, Rule Switching, and Response Reversal. Encyclopedia of Psychopharmacology, 1–7. 10.1007/978-3-642-27772-6_340-2

Burke, K. A., Takahashi, Y. K., Correll, J., Leon Brown, P., & Schoenbaum, G. (2009). Orbitofrontal inactivation impairs reversal of Pavlovian learning by interfering with “disinhibition” of responding for previously unrewarded cues. The European Journal of Neuroscience, 30(10), 1941–1946. 10.1111/J.1460-9568.2009.06992.X

Cadena-Valencia, J., García-Garibay, O., Merchant, H., Jazayeri, M., & de Lafuente, V. (2018). Entrainment and maintenance of an internal metronome in supplementary motor area. ELife, 7. 10.7554/ELIFE.38983

Cazettes, F., Mazzucato, L., Murakami, M., Morais, J. P., Augusto, E., Renart, A., & Mainen, Z. F. (2023). A reservoir of foraging decision variables in the mouse brain. Nature Neuroscience, 26(5), 840–849. 10.1038/S41593-023-01305-8

Chen, T.-W., Wardill, T. J., Sun, Y., Pulver, S. R., Renninger, S. L., Baohan, A., Schreiter, E. R., Kerr, R. A., Orger, M. B., Jayaraman, V., Looger, L. L., Svoboda, K., & Kim, D. S. (2013). Ultrasensitive fluorescent proteins for imaging neuronal activity. Nature, 499(7458), 295–300. 10.1038/nature12354

Dayan, P., & Daw, N. D. (2008). Decision theory, reinforcement learning, and the brain. In *Cognitive*, Affective and Behavioral Neuroscience (Vol. 8, Issue 4, pp. 429–453). Biblioteca Ayacucho. 10.3758/CABN.8.4.429

Ebbesen, C. L., & Brecht, M. (2017). Motor cortex - To act or not to act? In Nature Reviews Neuroscience (Vol. 18, Issue 11, pp. 694–705). Nature Publishing Group. 10.1038/nrn.2017.119

Ebbesen, C. L., Insanally, M. N., Kopec, C. D., Murakami, M., Saiki, A., & Erlich, J. C. (2018). More than just a “motor”: Recent surprises from the frontal cortex. Journal of Neuroscience, 38(44), 9402–9413. 10.1523/JNEUROSCI.1671-18.2018

Economo, M. N., Viswanathan, S., Tasic, B., Bas, E., Winnubst, J., Menon, V., Graybuck, L. T., Nguyen, T. N., Smith, K. A., Yao, Z., Wang, L., Gerfen, C. R., Chandrashekar, J., Zeng, H., Looger, L. L., & Svoboda, K. (2018). Distinct descending motor cortex pathways and their roles in movement. Nature, 563(7729), 79–84. 10.1038/s41586-018-0642-9

Erlich, J. C., Bialek, M., & Brody, C. D. (2011). A cortical substrate for memory-guided orienting in the rat. Neuron, 72(2), 330–343. 10.1016/j.neuron.2011.07.010

Everitt, B. J., & Robbins, T. W. (2016). Drug Addiction: Updating Actions to Habits to Compulsions Ten Years On. Annual Review of Psychology, 67, 23–50. 10.1146/ANNUREV-PSYCH-122414-033457

Floresco, S. B., Block, A. E., & Tse, M. T. L. (2008). Inactivation of the medial prefrontal cortex of the rat impairs strategy set-shifting, but not reversal learning, using a novel, automated procedure. Behavioural Brain Research, 190(1), 85–96. 10.1016/J.BBR.2008.02.008

Floresco, S. B., & Ghods-Sharifi, S. (2007). Amygdala-prefrontal cortical circuitry regulates effort-based decision making. Cerebral Cortex, 17(2), 251–260. 10.1093/cercor/bhj143

Gambino, F., Pagès, S., Kehayas, V., Baptista, D., Tatti, R., Carleton, A., & Holtmaat, A. (2014). Sensory-evoked LTP driven by dendritic plateau potentials in vivo. Nature, 515(7525), 116–119. 10.1038/nature13664

Gilad, A., Gallero-Salas, Y., Groos, D., & Helmchen, F. (2018). Behavioral Strategy Determines Frontal or Posterior Location of Short-Term Memory in Neocortex. Neuron, 99(4), 814–828.e7. 10.1016/J.NEURON.2018.07.029

Gremel, C. M., & Costa, R. M. (2013). Orbitofrontal and striatal circuits dynamically encode the shift between goal-directed and habitual actions. Nature Communications, 4, 2264. 10.1038/ncomms3264

Guo, Z. V., Li, N., Huber, D., Ophir, E., Gutnisky, D., Ting, J. T., Feng, G., & Svoboda, K. (2014). Flow of cortical activity underlying a tactile decision in mice. Neuron, 81(1), 179–194. 10.1016/j.neuron.2013.10.020

Haddon, J. E., & Killcross, S. (2006). Prefrontal Cortex Lesions Disrupt the Contextual Control of Response Conflict. Journal of Neuroscience, 26(11), 2933–2940. 10.1523/JNEUROSCI.3243-05.2006

Hattori, R., Danskin, B., Babic, Z., Mlynaryk, N., & Komiyama, T. (2019). Area-Specificity and Plasticity of History-Dependent Value Coding During Learning. Cell, 177(7), 1858–1872.e15. 10.1016/j.cell.2019.04.027

Hattori, R., & Komiyama, T. (2022). Context-dependent persistency as a coding mechanism for robust and widely distributed value coding. Neuron, 110(3), 502–515.e11. 10.1016/J.NEURON.2021.11.001

Hirokawa, J., Vaughan, A., Masset, P., Ott, T., & Kepecs, A. (2019). Frontal cortex neuron types categorically encode single decision variables. Nature, 576(7787), 446–451. 10.1038/S41586-019-1816-9

Inagaki, H. K., Chen, S., Daie, K., Finkelstein, A., Fontolan, L., Romani, S., & Svoboda, K. (2022). Neural Algorithms and Circuits for Motor Planning. Annual Review of Neuroscience, 45, 249–271. 10.1146/ANNUREV-NEURO-092021-121730

Inagaki, H. K., Chen, S., Ridder, M. C., Sah, P., Li, N., Yang, Z., Hasanbegovic, H., Gao, Z., Gerfen, C. R., & Svoboda, K. (2022). A midbrain-thalamus-cortex circuit reorganizes cortical dynamics to initiate movement. Cell, 185(6), 1065–1081.e23. 10.1016/J.CELL.2022.02.006

Ito, M., & Doya, K. (2009). Validation of decision-making models and analysis of decision variables in the rat basal ganglia. The Journal of Neuroscience: The Official Journal of the Society for Neuroscience, 29(31), 9861–9874. 10.1523/JNEUROSCI.6157-08.2009

Izquierdo, A., & Jentsch, J. D. (2012). Reversal learning as a measure of impulsive and compulsive behavior in addictions. Psychopharmacology, 219(2), 607–620. 10.1007/S00213-011-2579-7

Katahira, K., Matsuda, Y. T., Fujimura, T., Ueno, K., Asamizuya, T., Suzuki, C., Cheng, K., Okanoya, K., & Okada, M. (2015). Neural basis of decision making guided by emotional outcomes. Journal of Neurophysiology, 113(9), 3056–3068. 10.1152/JN.00564.2014

Kawai, R., Markman, T., Poddar, R., Ko, R., Fantana, A. L., Dhawale, A. K., Kampff, A. R., & Ölveczky, B. P. (2015). Motor Cortex Is Required for Learning but Not for Executing a Motor Skill. Neuron, 86(3), 800–812. 10.1016/j.neuron.2015.03.024

Kobak, D., Brendel, W., Constantinidis, C., Feierstein, C. E., Kepecs, A., Mainen, Z. F., Qi, X. L., Romo, R., Uchida, N., & Machens, C. K. (2016). Demixed principal component analysis of neural population data. ELife, 5(APRIL2016). 10.7554/ELIFE.10989

Komiyama, T., Sato, T. R., O’Connor, D. H., Zhang, Y.-X., Huber, D., Hooks, B. M., Gabitto, M., Svoboda, K., O?Connor, D. H., Zhang, Y.-X., Huber, D., Hooks, B. M., Gabitto, M., Svoboda, K., O’Connor, D. H., Zhang, Y.-X., Huber, D., Hooks, B. M., Gabitto, M., … Svoboda, K. (2010). Learning-related fine-scale specificity imaged in motor cortex circuits of behaving mice. Nature, 464(7292), 1182–1186. 10.1038/nature08897

Kopec, C. D., Erlich, J. C., Brunton, B. W., Deisseroth, K., & Brody, C. D. (2015). Cortical and Subcortical Contributions to Short-Term Memory for Orienting Movements. Neuron, 88(2), 367–377. 10.1016/j.neuron.2015.08.033

Krajbich, I., Armel, C., & Rangel, A. (2010). Visual fixations and the computation and comparison of value in simple choice. Nature Neuroscience, 13(10), 1292–1298. 10.1038/NN.2635

Maia, T. V, & Frank, M. J. (2011). From reinforcement learning models of the basal ganglia to the pathophysiology of psychiatric and neurological disorders. Nature Neuroscience, 14(2), 154–162. 10.1038/nn.2723.

Mathis, A., Mamidanna, P., Cury, K. M., Abe, T., Murthy, V. N., Mathis, M. W., & Bethge, M. (2018). DeepLabCut: markerless pose estimation of user-defined body parts with deep learning. Nature Neuroscience, 21(9), 1281–1289. 10.1038/S41593-018-0209-Y

Miller, P., & Katz, D. B. (2010). Stochastic transitions between neural states in taste processing and decision-making. Journal of Neuroscience, 30(7), 2559–2570. 10.1523/JNEUROSCI.3047-09.2010

Morris, R. W., Dezfouli, A., Griffiths, K. R., & Balleine, B. W. (2014). Action-value comparisons in the dorsolateral prefrontal cortex control choice between goal-directed actions. Nature Communications, 5, 4390. 10.1038/ncomms5390

Murakami, M., Shteingart, H., Loewenstein, Y., & Mainen, Z. F. (2017). Distinct Sources of Deterministic and Stochastic Components of Action Timing Decisions in Rodent Frontal Cortex. Neuron, 94(4), 908–919.e7. 10.1016/j.neuron.2017.04.040

Murakami, M., Vicente, M. I., Costa, G. M., & Mainen, Z. F. (2014). Neural antecedents of self-initiated actions in secondary motor cortex. Nature Neuroscience, 17(11), 1574–1582. 10.1038/nn.3826

Ostlund, S. B., Winterbauer, N. E., & Balleine, B. W. (2009). Evidence of action sequence chunking in goal-directed instrumental conditioning and its dependence on the dorsomedial prefrontal cortex. The Journal of Neuroscience: The Official Journal of the Society for Neuroscience, 29(25), 8280–8287. 10.1523/JNEUROSCI.1176-09.2009

Pachitariu, M., Stringer, C., Schröder, S., Dipoppa, M., Rossi, L. F., Carandini, M., & Harris, K. D. (2016). Suite2p: beyond 10,000 neurons with standard two-photon microscopy. BioRxiv, 061507. 10.1101/061507

Padoa-Schioppa, C. (2013). Neuronal origins of choice variability in economic decisions. Neuron, 80(5), 1322–1336. 10.1016/J.NEURON.2013.09.013

Padoa-Schioppa, C., & Assad, J. A. (2006). Neurons in the orbitofrontal cortex encode economic value. Nature, 441(7090), 223–226. 10.1038/NATURE04676

Passecker, J., Mikus, N., Malagon-Vina, H., Anner, P., Dimidschstein, J., Fishell, G., Dorffner, G., & Klausberger, T. (2019). Activity of Prefrontal Neurons Predict Future Choices during Gambling. Neuron, 101(1), 152–164.e7. 10.1016/J.NEURON.2018.10.050

Reep, R. L., Corwin, J. V., Hashimoto, A., & Watson, R. T. (1987). Efferent connections of the rostral portion of medial agranular cortex in rats. Brain Research Bulletin, 19(2), 203–221. 10.1016/0361-9230(87)90086-4

Reep, R. L., Goodwin, G. S., & Corwin, J. V. (1990). Topographic organization in the corticocortical connections of medial agranular cortex in rats. The Journal of Comparative Neurology, 294(2), 262–280. 10.1002/CNE.902940210

Rich, E. L., & Shapiro, M. (2009). Rat Prefrontal Cortical Neurons Selectively Code Strategy Switches. Journal of Neuroscience, 29(22), 7208–7219. 10.1523/JNEUROSCI.6068-08.2009

Rich, E. L., & Wallis, J. D. (2016). Decoding subjective decisions from orbitofrontal cortex. Nature Neuroscience, 19(7), 973–980. 10.1038/NN.4320

Rigotti, M., Barak, O., Warden, M. R., Wang, X. J., Daw, N. D., Miller, E. K., & Fusi, S. (2013). The importance of mixed selectivity in complex cognitive tasks. Nature, 497(7451), 585–590. 10.1038/NATURE12160

Russo, A. A., Bittner, S. R., Perkins, S. M., Seely, J. S., London, B. M., Lara, A. H., Miri, A., Marshall, N. J., Kohn, A., Jessell, T. M., Abbott, L. F., Cunningham, J. P., & Churchland, M. M. (2018). Motor Cortex Embeds Muscle-like Commands in an Untangled Population Response. Neuron, 97(4), 953–966.e8. 10.1016/J.NEURON.2018.01.004

Russo, A. A., Khajeh, R., Bittner, S. R., Perkins, S. M., Cunningham, J. P., Abbott, L. F., & Churchland, M. M. (2020). Neural Trajectories in the Supplementary Motor Area and Motor Cortex Exhibit Distinct Geometries, Compatible with Different Classes of Computation. Neuron, 107(4), 745–758.e6. 10.1016/J.NEURON.2020.05.020

Schoenbaum, G., Nugent, S. L., Saddoris, M. P., & Setlow, B. (2002). Orbitofrontal lesions in rats impair reversal but not acquisition of go, no-go odor discriminations. Neuroreport, 13(6), 885–890. 10.1097/00001756-200205070-00030

Siniscalchi, M. J., Phoumthipphavong, V., Ali, F., Lozano, M., & Kwan, A. C. (2016). Fast and slow transitions in frontal ensemble activity during flexible sensorimotor behavior. Nature Neuroscience, 19(9), 1234–1242. 10.1038/nn.4342

Soltani, A., & Izquierdo, A. (2019). Adaptive learning under expected and unexpected uncertainty. Nature Reviews. Neuroscience, 20(10), 635–644. 10.1038/S41583-019-0180-Y

Sul, J. H., Jo, S., Lee, D., & Jung, M. W. (2011). Role of rodent secondary motor cortex in value-based action selection. Nature Neuroscience, 14(9), 1202–1208. 10.1038/nn.2881

Sul, J. H., Kim, H., Huh, N., Lee, D., & Jung, M. W. (2010). Distinct roles of rodent orbitofrontal and medial prefrontal cortex in decision making. Neuron, 66(3), 449–460. 10.1016/j.neuron.2010.03.033

Sutton, R. S., & Barto, A. G. (2018). Reinforcement Learning: An Introduction (Adaptive Computation and Machine Learning). The MIT Press 2018. https://mitpress.mit.edu/books/reinforcement-learning-second-edition

Svoboda, K., & Li, N. (2018). Neural mechanisms of movement planning: motor cortex and beyond. In Current Opinion in Neurobiology (Vol. 49, pp. 33–41). Elsevier Ltd. 10.1016/j.conb.2017.10.023

Tombaz, T., Dunn, B. A., Hovde, K., Cubero, R. J., Mimica, B., Mamidanna, P., Roudi, Y., & Whitlock, J. R. (2020). Action representation in the mouse parieto-frontal network. Scientific Reports, 10(1). 10.1038/S41598-020-62089-6

Tsutsui, K. I., Grabenhorst, F., Kobayashi, S., & Schultz, W. (2016). A dynamic code for economic object valuation in prefrontal cortex neurons. Nature Communications, 7. 10.1038/NCOMMS12554

Turner, R. S., & Desmurget, M. (2010). Basal ganglia contributions to motor control: a vigorous tutor. Current Opinion in Neurobiology, 20(6), 704–716. 10.1016/J.CONB.2010.08.022

Waltz, J. A., & Gold, J. M. (2007). Probabilistic reversal learning impairments in schizophrenia: further evidence of orbitofrontal dysfunction. Schizophrenia Research, 93(1–3), 296–303. 10.1016/J.SCHRES.2007.03.010

Wang, A. Y., Miura, K., & Uchida, N. (2013). The dorsomedial striatum encodes net expected return, critical for energizing performance vigor. Nature Neuroscience, 16(5), 639–647. 10.1038/NN.3377

Wang, X. J. (2008). Decision Making in Recurrent Neuronal Circuits. In Neuron (Vol. 60, Issue 2, pp. 215–234). Neuron. 10.1016/j.neuron.2008.09.034

Wardak, C., Ramanoël, S., Guipponi, O., Boulinguez, P., & Ben Hamed, S. B. (2012). Proactive inhibitory control varies with task context. The European Journal of Neuroscience, 36(11), 3568–3579. 10.1111/J.1460-9568.2012.08264.X

Wilson, R. C., Takahashi, Y. K., Schoenbaum, G., & Niv, Y. (2014). Orbitofrontal cortex as a cognitive map of task space. Neuron, 81(2), 267–279. 10.1016/J.NEURON.2013.11.005

Yang, J. H., & Kwan, A. C. (2021). Secondary motor cortex: Broadcasting and biasing animal’s decisions through long-range circuits. International Review of Neurobiology, 158, 443–470. 10.1016/BS.IRN.2020.11.008

Yang, W., Kanodia, H., & Arber, S. (2023). Structural and functional map for forelimb movement phases between cortex and medulla. Cell, 186(1), 162–177.e18. 10.1016/J.CELL.2022.12.009

