## Supplemental Figures for "Secondary motor cortex tracks decision value and supports behavioral flexibility during non-instructed choice"

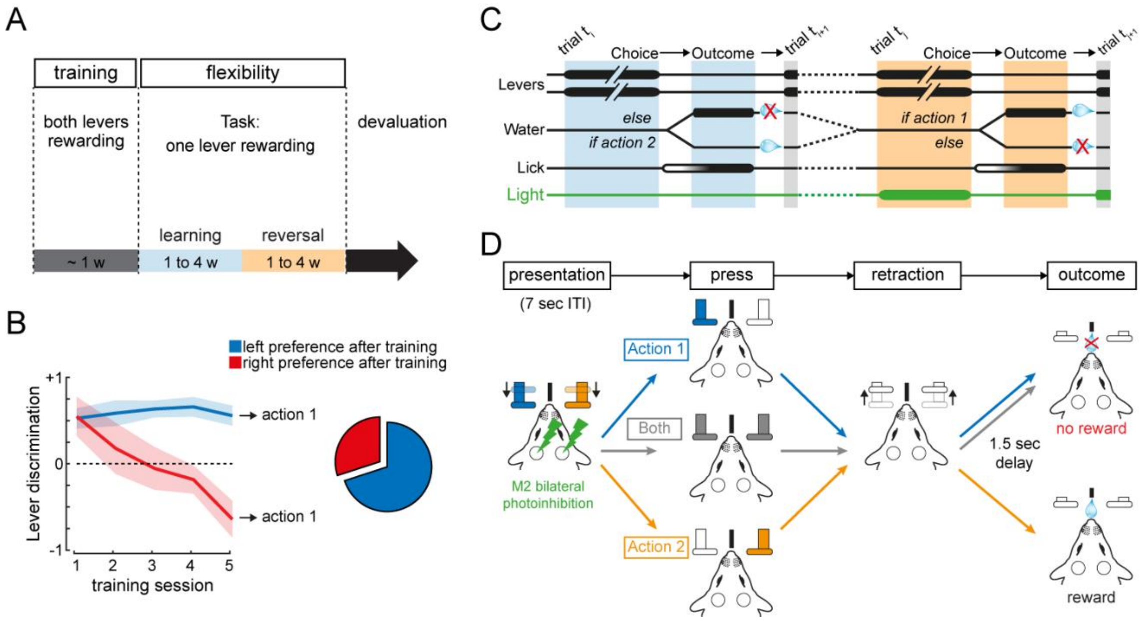


**Supp Figure 1: Reinforcement learning task. A)** Time flow of the behavior task. **B)** Bias of the mice after training. **C)** Schematic representation of a trial from the learning cycle (blue) and the reversal cycle with M2 illumination (orange). **D)** Events in a trial from reversal cycle.

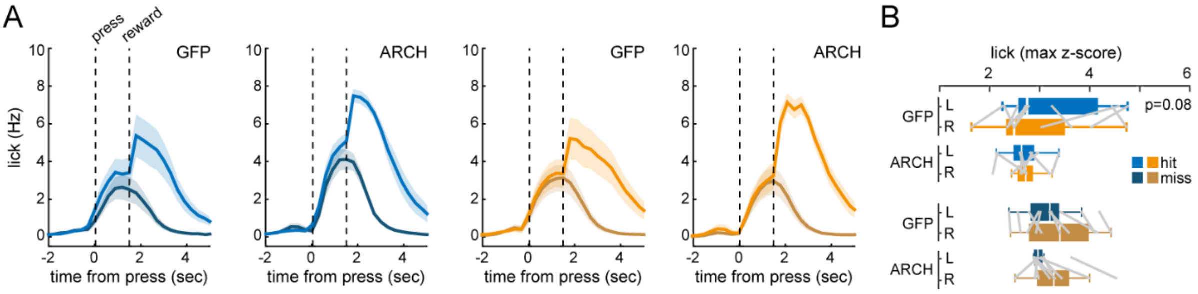


**Supp Figure 2: Lick frequency during the different trials. A)** Lick frequency throughout the hit (light) and miss (dark) trials. Blue from learning cycle, orange from reversal cycle. Shaded bars are sem. **B)** Max z-score of the lick on the different trials. Boxplots represent mean and interquartile range. Error bars are sem.

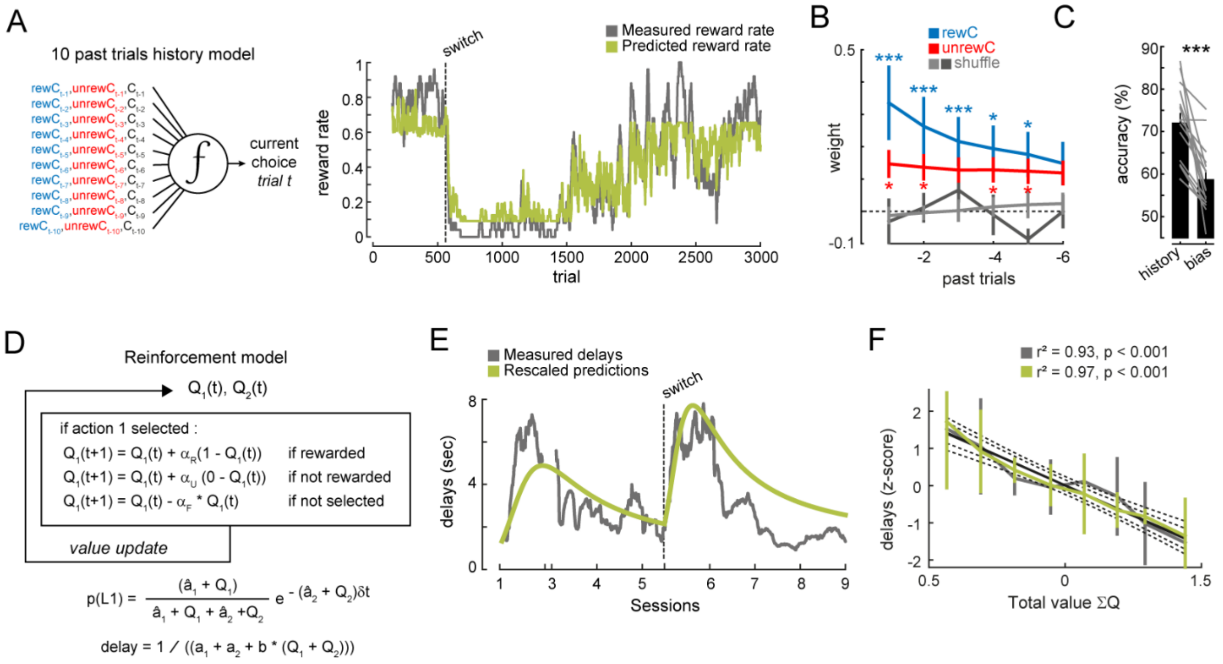


**Supp Figure 3: Reinforcement learning model based on trials history. A)** *Left panel*: Illustration of the regression model based on the history of the outcome of the 10 past trials for predicting the current choice. *Right panel*: Representation of the predicted reward rate by the model (green) and measured (gray). **B)** Weight of the regression model predicting choice having in account the different past trials (blue, past rewarded choices; red, past unrewarded choices; light grey, shuffled data from rewarded trials; dark grey, shuffled data from unrewarded trials). Only the regression weights for the past 6 trials are shown. **C)** Comparison between the regression model based on the history of the outcome and the model based on a constant term that accounts for by the bias for the levers. **D)** Illustration of the reinforcement learning (RL) model. **E)** Representation of the predicted delays by the RL-model (green) and the observed delays (grey), during learning and reversal cycles. Dashed black line marks the switch between cycles. **F)** Correlation between the press delays predicted by the RL-model (green) and measured (grey) and the total value (ΣQ) obtained by the RL-model.

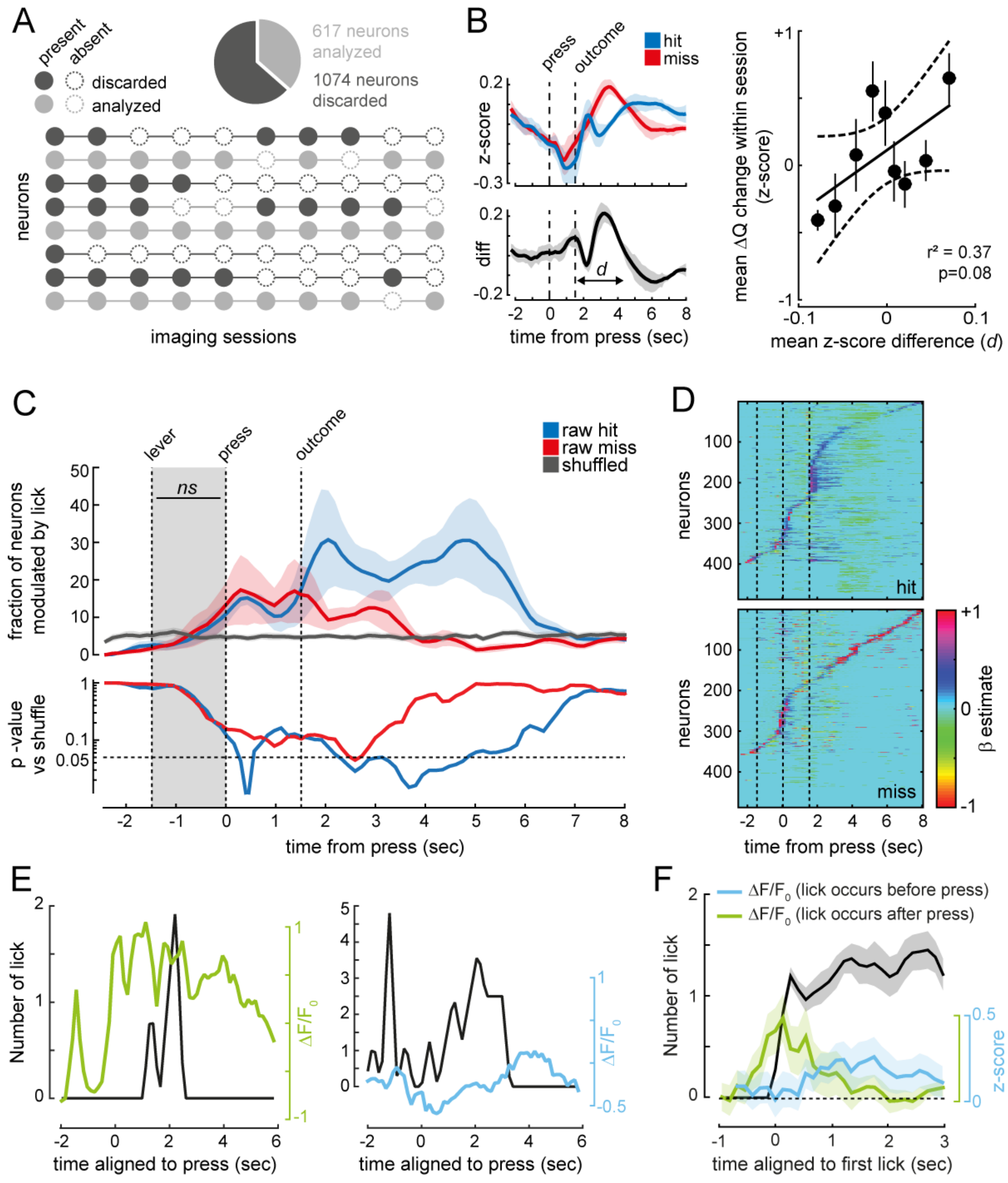


**Supp Figure 4: M2 neuronal activity modulation by lick. A)** Illustration of the criteria to select neurons for further analysis. **B)** *Top left:* $\Delta F/F_0$ (z-score) throughout the trial period in hit (reward) and miss (unrewarded) trials. *Bottom left:* the difference in $\Delta F/F_0$ signals between miss and hit trials within session (*d*) was averaged across all sessions. *Right:* correlation between "d" and the increase in ΔQ that occurs between the beginning and end of each session. **C)** *Top panel*: Fraction of neurons in M2 that are modulated by lick throughout the trial in hit (blue trace) and miss (red trace) trials and shuffled data (gray trace) using a multiple linear regression. *Bottom panel*: p-value of the calculated vs. shuffled data. **D)** β coefficients of the linear regression between $\Delta F/F_0$ and the number of Licks for hit (*top*) and miss (*bottom*) trials throughout the different epochs of the trial. **E)** *Left panel*: Number of licks (black) and $\Delta F/F_0$ (green) in a trial in which the first lick occurs after press. *Right panel*: Number of licks (black) and $\Delta F/F_0$ (blue) in a trial in which the first lick occurs before press. **F)** Mean number of licks (black) and mean $\Delta F/F_0$ in the trials in which the first lick occurs after press (green) or before press (blue) aligned to the first lick. Shaded bars are sem.

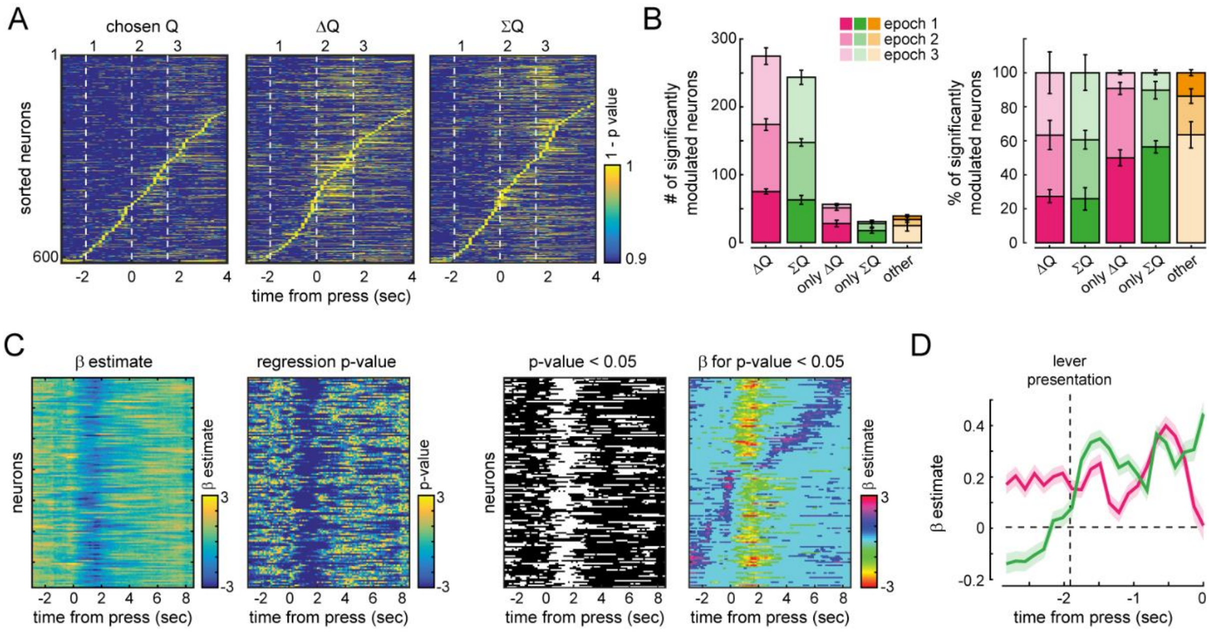


**Supp Figure 5: M2 neuronal activity modulation by ΔQ**. **A)** *p*-value associated with its β regression coefficient for three different regressors – chosen Q (value of the chosen action), ΔQ and ∑Q, for all sorted neurons, during the three different epochs of a trial. **B)** *Left panel:* Number of significantly modulated neurons for the regressors ΔQ and ∑Q in the three different epochs of the trials (1, during lever presentation; 2, between press and outcome; 3, after outcome or inter-trial interval). ΔQ, all neurons modulated by ΔQ; ∑Q, all neurons modulated by ∑Q; only ΔQ, all neurons modulated only by ΔQ and no other regressors; only ∑Q, all modulated neurons only by ∑Q and no other regressors; other, neurons modulated by other than ΔQ and ∑Q regressors. *Right panel*: Percentage of significantly modulated neurons for the different regressors in the three different epochs of the trials. **C)** Illustration of the method based on linear regression used to extract significantly modulated neurons. *Left panel:* illustrative β coefficients from the regression model during a trial from all sorted neurons. *Second panel: p*-value of the regression during a trial. *Third panel: p*-value < 0.05 of the regression during a trial. *Right panel:* β coefficients for *p*-value < 0.05 during a trial. **D)** Dynamic of β coefficients during a trial from neurons positively or negatively modulated by ΔQ (pink) and ∑Q (green).

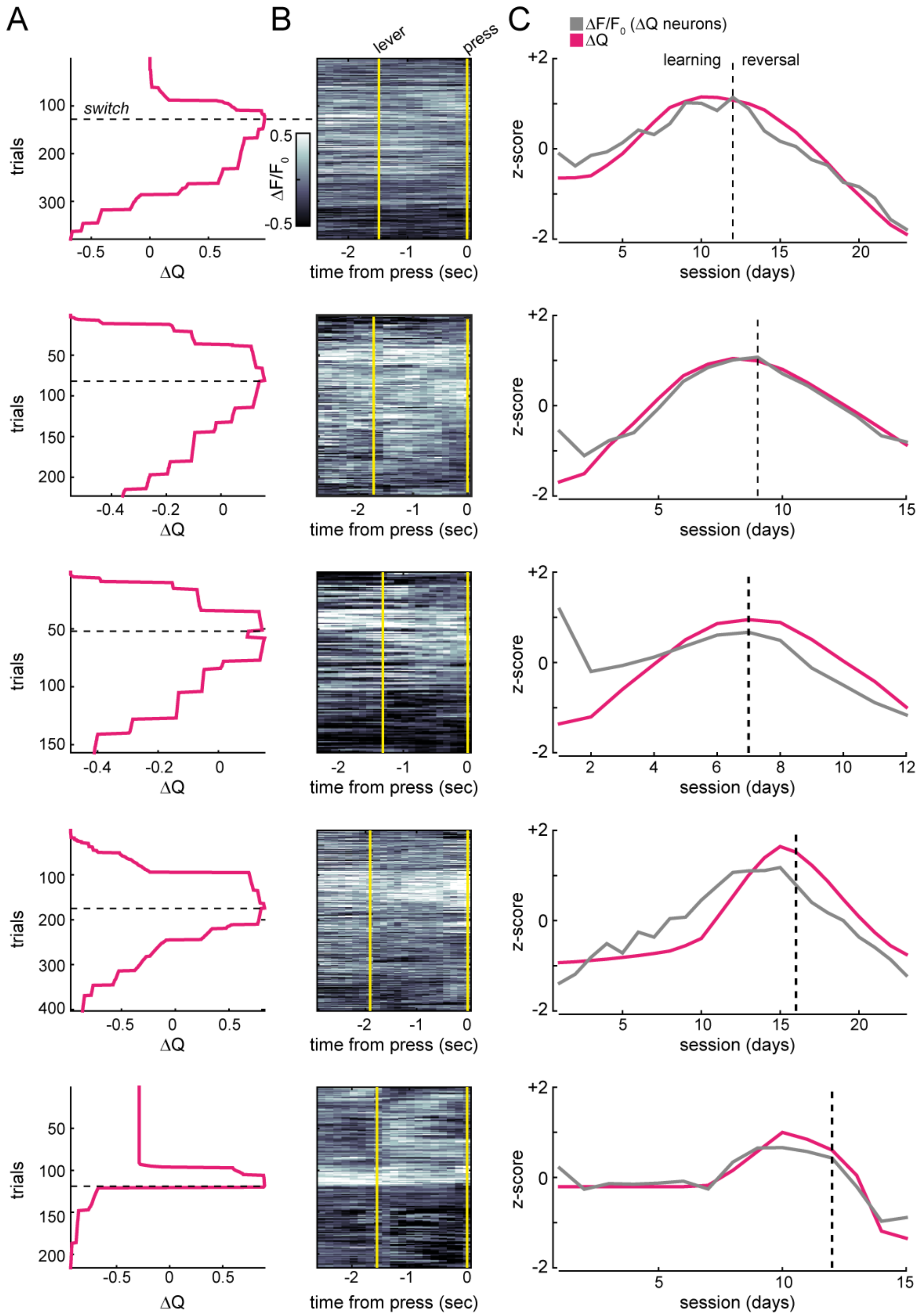


**Supp Figure 6: Monotic representation of decision variable in the M2 of 5 individual mice. A)** Dynamic of ΔQ across trials. **B)** Heat map of ΔF/F averaged across ΔQ-modulated neurons. **C)** z-score of ΔF/F (averaged for each session) of ΔQ-modulated neurons (grey) and of ΔQ (pink, averaged for each session) measured between lever presentation and lever-pressing, during learning and reversal, separated by the dashed black line.

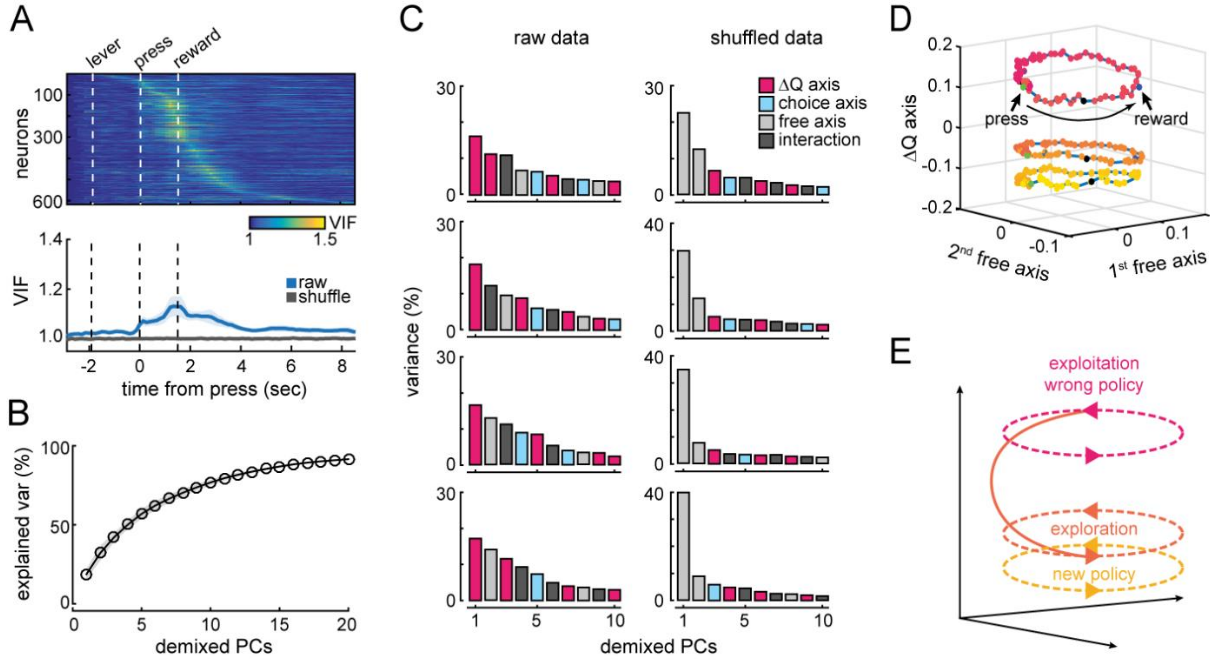


**Supp Figure 7: demixed PCA. A)** Variance inflation factor (VIF) for ΔQ from all sorted neurons during the different epoch of a trial. VIF lower than 2 indicates that multicollinearity between regressors (see methods) did not occur. **B)** Explained variance by the different number of demixed PC. **C)** Examples from 4 different other mice than in Figure 6 of the distribution of the variance in the first 10 PCs in task-dependent and task-independent dimensions, in the raw data (*left*) and the shuffled data (*right*). **D)** The same data set than in Figure 6 is projected onto the ΔQ axis and the two task-independent axes with the greatest variance. **E)** Schematic representation of the trajectories represented on panel c (dashed lines) and hypothetical transition from one to the other while ΔQ decreased during reversal cycle.
